# A cell type-specific mechanism driving the rapid antidepressant effects of transcranial magnetic stimulation

**DOI:** 10.1101/2025.01.29.635537

**Authors:** Michael W. Gongwer, Alex Qi, Alexander S. Enos, Sophia A. Rueda Mora, Cassandra B. Klune, Meelan Shari, Adrienne Q. Kashay, Owen H. Williams, Aliza Hacking, Jack P. Riley, Gary A. Wilke, Yihong Yang, Hanbing Lu, Andrew F. Leuchter, Laura A. DeNardo, Scott A. Wilke

## Abstract

Repetitive transcranial magnetic stimulation (rTMS) is an emerging treatment for brain disorders, but its therapeutic mechanism is unknown. We developed a novel mouse model of rTMS with superior clinical face validity and investigated the neural mechanism by which accelerated intermittent theta burst stimulation (aiTBS) – the first rapid-acting rTMS antidepressant protocol – reversed chronic stress-induced behavioral deficits. Using fiber photometry, we showed that aiTBS drives distinct patterns of neural activity in intratelencephalic (IT) and pyramidal tract (PT) projecting neurons in dorsomedial prefrontal cortex (dmPFC). However, only IT neurons exhibited persistently increased activity during both aiTBS and subsequent depression-related behaviors. Similarly, aiTBS reversed stress-related loss of dendritic spines on IT, but not PT neurons, further demonstrating cell type-specific effects of stimulation. Finally, chemogenetic inhibition of dmPFC IT neurons during rTMS blocked the antidepressant-like behavioral effects of aiTBS. Thus, we demonstrate a prefrontal mechanism linking rapid aiTBS-driven therapeutic effects to cell type-specific circuit plasticity.

## Introduction

Brain stimulation promises to revolutionize treatment of brain disorders characterized by aberrant circuit function^1^. Transcranial magnetic stimulation (TMS) is a non-invasive form of focal neuromodulation that drives neural activity in a target region using electromagnetic pulses^2–4^. TMS is the dominant form of clinical brain stimulation used to treat depression and a number of other disorders^5–8^. When employed to treat neuropsychiatric disorders, repetitive TMS (rTMS) pulse sequences are used to produce lasting changes in brain function^9,10^. While this is often effective, many patients fail to respond or have residual symptoms^11,12^, and several brain disorders do not have approved rTMS-based therapies. The lack of a mechanistic understanding of rTMS impedes the rational optimization of protocols that target dysfunctional circuits underlying specific symptoms or disorders.

Prefrontal cortex (PFC) dysfunction is a hallmark of major depressive disorder^13,14^. In clinical rTMS, the coil is targeted to specific cortical regions based on theories about the brain networks disrupted in a particular disorder^15–17^. While work from preclinical models has suggested that rTMS may drive plasticity in specific cortical cell types^18–22^, no studies to date have identified causal mechanisms driving changes in behavior. As a result, it remains unclear whether rTMS drives lasting therapeutic effects by modifying specific cell types or circuits within the clinical target region *in vivo*. Progress towards discovering these circuit mechanisms has been limited by a lack of preclinical animal models of rTMS with strong face validity for how rTMS is delivered clinically. Recently, a prefrontal-targeted, accelerated intermittent theta burst stimulation (aiTBS) rTMS protocol, which compresses a typical 6-week treatment course to just 5 days, has been shown to rapidly reverse symptoms of depression and produce long-lasting effects^23–25^. Here we developed a novel preclinical model of rTMS and used it to discover how aiTBS drives cell-type specific plasticity in PFC *in vivo*.

## Results

### Establishing a novel preclinical rodent model of rTMS

In order to study the circuit mechanisms underlying the therapeutic effects of rTMS, we needed a rodent model that allowed us to deliver clinically effective protocols and mimicked how patients are treated with TMS in the clinic. However, scaling rTMS coils to the smaller rodent brain poses significant challenges^26,27^. Prior preclinical models used anesthetized mice and oversized coils that likely stimulated most of the brain, or miniaturized coils that were too weak to elicit action potentials^28–33^. Our group recently developed and extensively validated a novel coil design that overcomes these limitations, enabling highly focal (<2mm), suprathreshold stimulation of a cortical subregion in the rodent brain^34–37^. Using this coil, we developed a system that enables us to combine delivery of clinical rTMS protocols with rigorous investigation of neural circuit mechanisms in the awake mouse (Fig. 1A). Our model mimics how TMS patients are treated in the clinic, with the head held stable and in an awake state. Indeed, rTMS delivered under anesthesia can have effects that are opposite of those observed in awake animals, underscoring the translational importance of brain state ^28^.

**Fig. 1.**
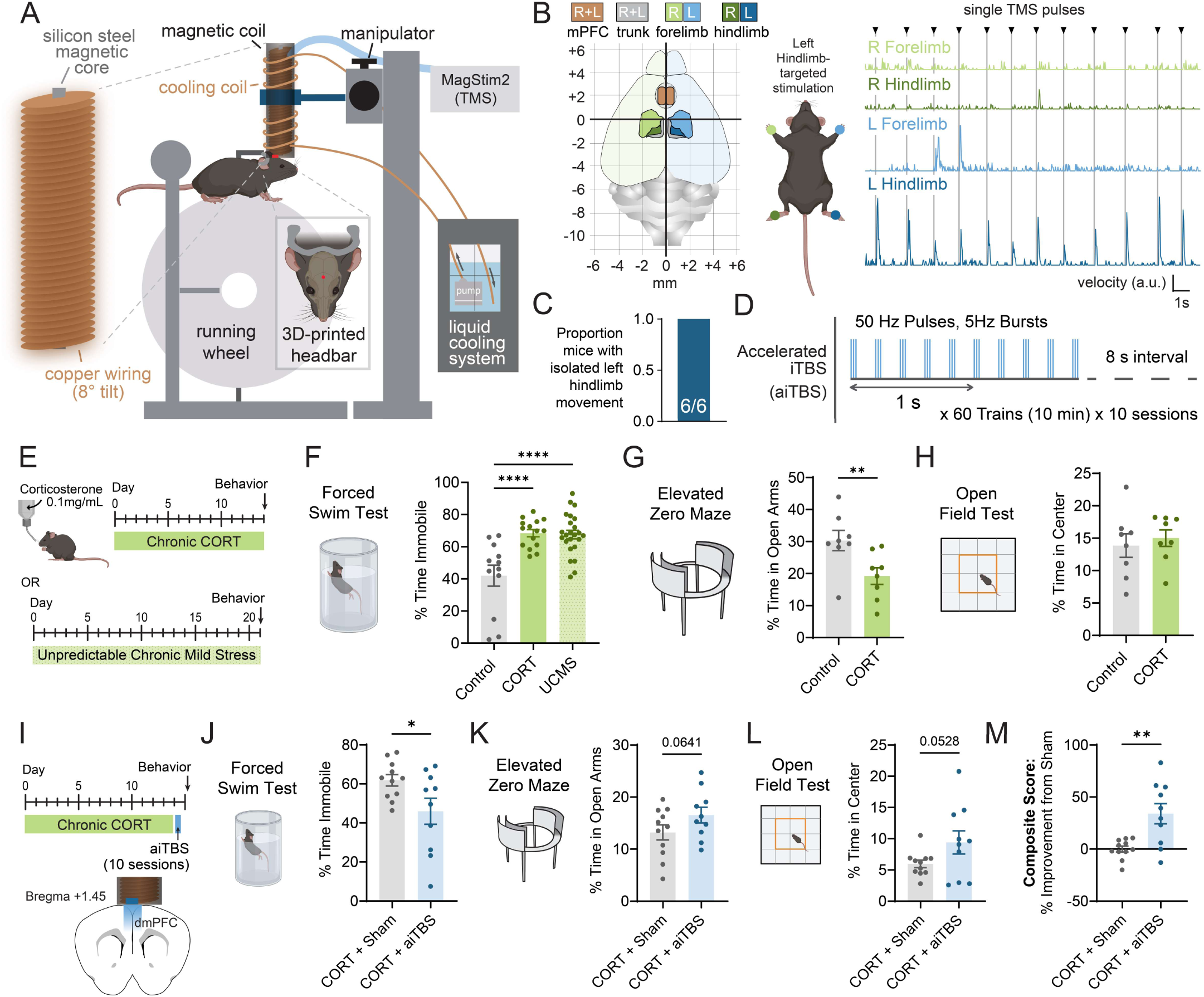
Focal aiTBS over dmPFC reverses behavioral effects of chronic stress. (**A**) Setup for rTMS treatment. (**B**) Magnet focality test. Left, map of limb representation in motor cortex. Right, limb velocity upon magnetic stimulation of left hindlimb region of motor cortex. (**C**) Proportion of mice showing isolated left hindlimb movements upon stimulation. n=6 mice. (**D**) Accelerated iTBS protocol. (**E**) Timeline of stress protocols used. (**F**) Effects of CORT and UCMS on immobility in the forced swim test (Control n=12, CORT n=15, UCMS n=24 mice; one-way ANOVA with Tukey’s multiple comparisons test). (**G,H**) Effects of CORT on anxiety-like behavior in the EZM and OFT (Control n=8, CORT n=8 mice; unpaired t-test). (**I**) Experimental strategy to assess behavioral effects of aiTBS treatment following chronic stress. (**J, K, L**) Effects of aiTBS treatment on behavior in FST, EZM, and OFT (CORT+Sham n=11, CORT+aiTBS n=10 mice; unpaired t-test). (**M**) Composite score of behavioral outcomes in FST, EZM, and OFT. Score was calculated as the average percent improvement from the mean of the sham-treated group across all assays (CORT+Sham n=11, CORT+aiTBS n=10 mice; unpaired t-test). Mouse diagrams were generated using Biorender.com. For detailed statistical analysis see Table S1. All error bars reflect mean ± SEM. *p<0.05, **p<0.01, ****p<0.0001.

We first verified the focality of stimulation in the awake mouse brain. To do so, we measured evoked motor responses when the coil hotspot – the location of the sharply focused electric field – was positioned over the hindlimb region of the primary motor cortex. Single pulses delivered to this region, which spans approximately 1 mm^2^ in the mouse brain^38,39^, elicited tightly time-locked motor responses limited to the contralateral hindlimb, confirming highly focal, suprathreshold stimulation (Fig. 1B,C). This is consistent with our previous reports, using the same coil, which used activity-dependent labeling and motor-evoked potentials to demonstrate similar focality^34,35^. To ensure precise and reliable positioning of the coil hotspot, we marked the region to be stimulated using stereotaxic coordinates. The TMS coil was mounted on a fine 3-axis manipulator to allow precise targeting in head-fixed mice that were freely running on a wheel (Fig. 1A).

While high-frequency clinical protocols are the most commonly used protocols in patients, they have been difficult to model in rodents because miniaturized coils tend to easily overheat^26^. To overcome this challenge, we surrounded our coil with a custom-designed, liquid-cooled heatsink, which allowed us to efficiently deliver longer clinical protocols (Fig. 1A). The coil was connected to a FDA-approved clinical stimulator capable of delivering theta burst protocols, which more efficiently elicit effects at lower stimulation intensities^40^. We utilized a recently developed aiTBS clinical protocol which targets the PFC to rapidly reverse depressive symptoms^23,24,41^. Our experiments used 1800 pulses of iTBS deployed as 10 sessions occurring once every hour, identical to the clinical protocol effective in patients (Fig. 1D)^23,24,41^. We delivered rTMS and sham treatment simultaneously, using identical side-by-side setups (Fig. S1).

### aiTBS reverses the behavioral effects of chronic stress

Depression is a highly pleiotropic condition driven by both environmental and genetic factors. Chronic stress is a major risk factor for depression and other psychiatric disorders^42,43^. Animal models of chronic stress reliably induce dendritic spine loss, dorsomedial PFC (dmPFC) hypofunction and other cellular and circuit disturbances^44–47^. These changes are thought to disrupt core functions of dmPFC including motivation, effort-reward valuation, and decision-making, manifesting as disordered behavior^46,47^. Patients suffering from depression often exhibit symptoms that are thought to arise from these deficits, including low resilience in the face of acute stressors^42,48^. Recently, interventions such as ketamine and psychedelic compounds have been shown to produce rapid-acting antidepressant behavioral effects by reversing the neurobiological disturbances generated by chronic stress^46,49,50^. We hypothesized that an accelerated rTMS protocol like aiTBS may elicit similar changes.

To better understand how rTMS drives antidepressant effects, we applied our novel rodent TMS system to a mouse model of chronic stress. We modeled the effects of stress using a well-validated protocol: chronic administration of the steroid hormone corticosterone (CORT; Fig. 1E)^46,47,51–53^. Then, using a series of behavioral tests, we examined how mice responded when exposed to acutely stressful situations. CORT reliably increased passive coping (immobility) in the forced swim test (FST), consistent with the decreased effortful persistence characteristic of a depression-like behavioral state (Fig. 1F). To ensure these behavioral effects were robust, we repeated these experiments using a different chronic stress protocol in which mice are exposed to a series of environmental stressors (unpredictable chronic mild stress; UCMS)^54,55^. Similar to CORT, mice that experienced UCMS exhibited increased immobility in the FST (Fig. 1F).

Depression is highly comorbid with anxiety, and rTMS can improve symptoms of both^56,57^. Therefore, we also examined how CORT impacted behavior in two commonly used assays for anxiety-like behaviors in rodents: the elevated zero maze (EZM) and open field test (OFT). The EZM and OFT measure how much time mice spend in exposed parts of an apparatus, where they are more vulnerable to predation. CORT decreased the time spent in the open arms of the EZM, suggesting an increase in anxiety-like behavior (Fig. 1G). We found no behavioral effects of CORT in the OFT, which has less physical distinction between the safe and vulnerable areas of the arena compared to EZM (Fig. 1H). Thus, chronic CORT produced a behavioral state with anxiety- and depression-like features, which we could use to interrogate the potential therapeutic mechanisms underlying rTMS treatment of depression.

We hypothesized that aiTBS targeting dmPFC would reverse behavioral deficits observed in our CORT model of chronic stress. To test this, mice were placed on CORT, treated with aiTBS (10 aiTBS sessions in 1 day), and then assessed for behavioral outcomes the following day (Fig. 1I). aiTBS treatment significantly reduced immobility in the FST compared to sham-treated controls (Fig. 1J), consistent with an antidepressant-like effect. We also observed strong trends toward increased time in the open arms of the EZM and in the center of the OFT following aiTBS treatment (Fig. 1K,L). To examine the overall response to aiTBS across these assays, we calculated a composite score based on behavior in the FST, EZM, and OFT. The results indicated that aiTBS significantly improved anxiety- and depression-like behaviors (Fig. 1M). We also replicated the effects of aiTBS on FST behavior in mice exposed to UCMS instead of CORT (Fig. S2). Thus, our model exhibits strong face validity for clinical rTMS and recapitulates the rapid therapeutic effects seen with accelerated rTMS for depression.

Chronic stress models of depression are also characterized by other behavioral features, including anhedonia and reduced effortful persistence^58–60^. We used additional assays to test whether our novel model of aiTBS could ameliorate such behavioral impairments. We used the sucrose preference test (SPT) to measure behavioral anhedonia^61^, a proposed corollary of the reduced ability to experience pleasure which is a core feature of depression^58^. Chronic CORT administration decreased sucrose preference from a pre-stress baseline, and aiTBS reversed this effect (Fig. S3A-C). As an additional measure of effortful behavioral persistence beyond the FST, we used the sinking platform test^60^. This assay measures the propensity to retain goal-directed actions under adverse conditions (Fig. S3D-E). Chronic CORT decreased behavioral persistence (measured as the number of platforms climbed), and aiTBS reversed this effect (Fig. S3F-H). These findings indicate that dmPFC-targeted aiTBS improves the behavioral consequences of chronic stress related to both anhedonia and behavioral persistence.

Clinical depression is frequently characterized by imbalances in the drive to approach rewarding stimuli and avoid aversive stimuli^62^, leading to maladaptive decision-making in the face of approach-avoidance conflicts^62^. We therefore evaluated the effects of aiTBS on stress-induced disruptions in the balance of reward approach and threat avoidance behaviors. We used an approach-avoidance conflict assay where mice must balance approaching a liquid reward with avoiding a cued footshock (Fig. S4A-C)^63^. Mice must balance approach and avoidance behaviors during periods of elevated threat (tone), or relative safety during inter-tone intervals (ITI). Across training, mice from all experimental groups displayed similar increases in avoidance and reductions in reward-seeking during the tone period (Fig. S4D). By the end of training and during a retrieval session without foot shocks, stressed, aiTBS-treated and non-stressed control mice had a significantly higher adaptive preference for the reward zone during the ITI compared to stressed, sham-treated mice (Fig. S4E-G). Therefore, aiTBS reversed the stress-induced decreases in reward-seeking behavior during approach-avoidance conflict. These changes were specific to intervals when the threatening cue was absent, when approach was adaptive and avoidance was maladaptive. Taken together, these findings demonstrate that a clinically effective aiTBS protocol can rapidly reverse multiple stress-induced behavioral deficits relevant to depression.

### aiTBS elicits cell type-specific neuromodulation

Having developed an experimentally tractable system for studying rTMS, we sought to discover the neural mechanisms underlying the therapeutic behavioral effects of aiTBS. Patients suffering from depression exhibit reduced motivation to exert effort, excessive avoidance of aversive stimuli and altered encoding of emotional stimuli in PFC, amongst other deficits^64^. Thus, effective treatment with aiTBS likely alters activity in the cell types and circuits that underlie these behavioral changes. Within dmPFC, projection-defined classes of neurons play distinct roles in behavior, and manipulating these projections can produce distinct effects in the FST and other depression-related behaviors^65–70^. The PFC contains intratelencephalic (IT) and pyramidal tract (PT) projection classes^71,72^, two non-overlapping excitatory cell types that are defined by their long-range projection targets and are highly conserved across species and between cortical regions^73,74^. IT neurons project to cortical and striatal targets, while PT neurons project to midbrain, hindbrain, striatal, and thalamic targets^73,74^. dmPFC IT and PT neurons receive markedly distinct local and long-range synaptic inputs and exhibit unique molecular profiles^75–79^. Growing evidence suggests that imbalanced IT/PT function in PFC may underlie neuropsychiatric disease states, including depression^80^. Thus, we hypothesized that IT and PT neurons may differentially respond to aiTBS and/or play distinguishable roles in the therapeutic effects of aiTBS. To test this, we recorded IT and PT neuronal activity during iTBS, followed by recording during effortful coping and encoding of appetitive and aversive stimuli.

We reasoned that if we could detect differential effects of aiTBS on IT and PT neuron activity, this would generate insight into the loci of rTMS-driven circuit plasticity. To investigate this, we first used fiber photometry to record activity of dmPFC IT and PT neurons expressing the genetically encoded calcium sensor GCaMP7f (Fig. 2A-C, S5). We developed a surgical approach to implant the optical fiber from the side of the brain, allowing simultaneous rTMS delivery and recording of neural activity (Fig. 2B). After mice received chronic CORT treatment, we measured how IT and PT neurons acutely responded during aiTBS (Fig. 2D).

**Fig. 2.**
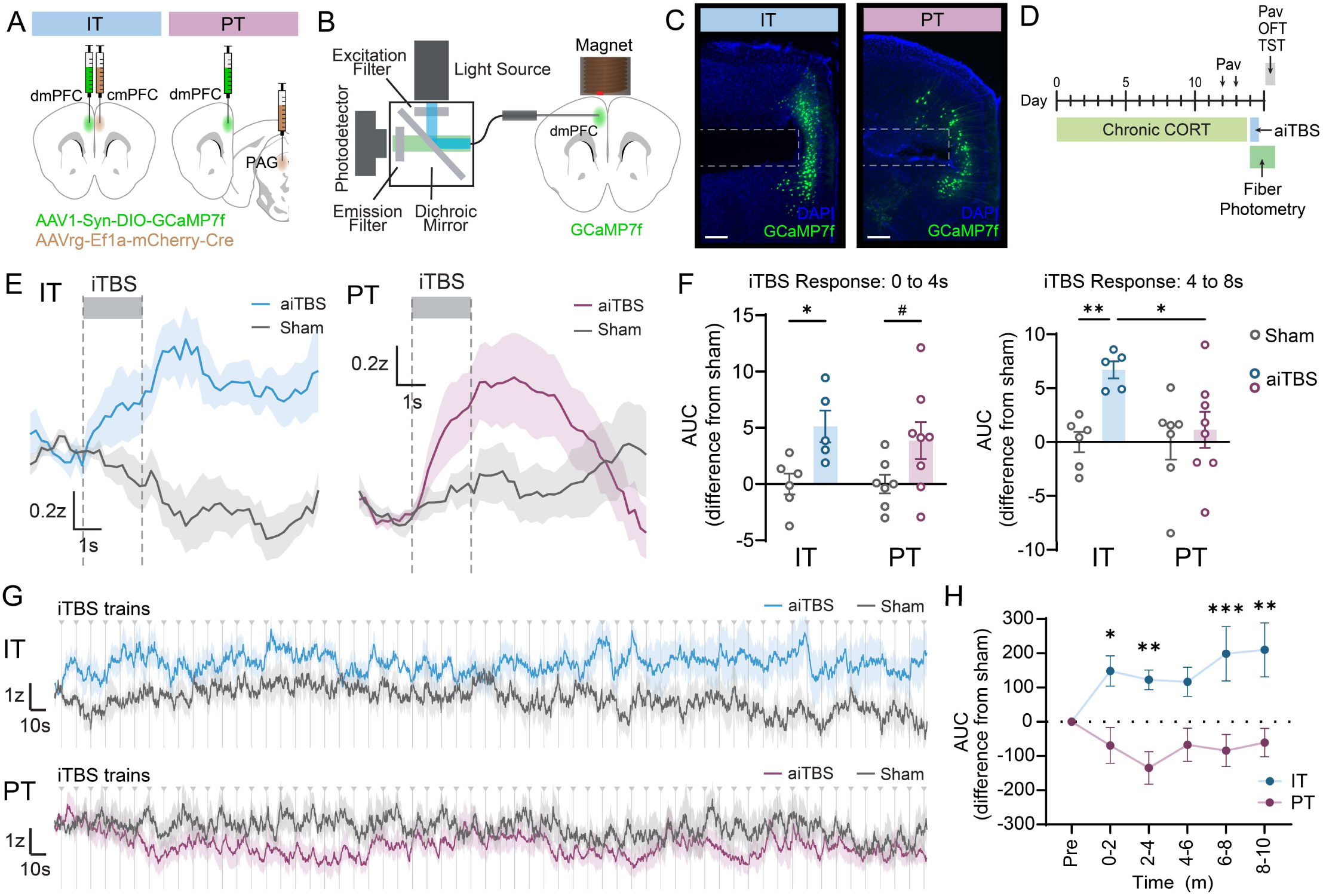
aiTBS drives cell-type specific neuromodulation. (**A**) Viral strategy to record neural activity in IT and PT neurons of left dmPFC. (**B**) Fiber photometry setup with lateral fiber implant to record cell-type specific neural activity during and after rTMS. (**C**) Representative images of fiber placement and viral expression to record from IT and PT neurons. Scale bars 200μm. (**D**) Experimental timeline. Mice were placed on CORT for 14 days. On days 12 and 13, mice underwent Pavlovian conditioning for rewarding and aversive stimuli. On day 14, mice received aiTBS treatment, and fiber photometry signals were recorded during the first session of aiTBS. On day 15, fiber photometry signals were recorded during Pavlovian stimuli, OFT, and TST. (**E**) Fiber photometry recordings from IT (left) and PT (right) neurons during individual aiTBS trains. (**F**) Area under the curve (AUC) values calculated from plots shown in E, normalized to sham-treated mice, from IT and PT neurons during time windows relative to the start of stimulation (IT sham n=6, IT aiTBS n=5, PT sham n=7, PT aiTBS n=8 mice; two-way ANOVA with Sidak’s multiple comparisons test). (**G**) IT and PT activity across the aiTBS session. (**H**) AUC values from different time bins across the session (IT sham n=6, IT aiTBS n=5, PT sham n=7, PT aiTBS n=8 mice; two-way ANOVA with Sidak’s multiple comparisons test). For detailed statistical analysis see Table S1. Lines and shading reflect mean ± SEM of average traces across mice. All error bars reflect mean ± SEM. #p<0.1, *p<0.05, **p<0.01, ***p<0.001.

For each mouse, we recorded GCaMP fluorescence (a proxy for neural activity) during the first 10-minute aiTBS session. When normalized to the pre-train period, iTBS elicited a ramping increase in activity for both IT and PT neurons, which persisted for at least 4 seconds into the intertrain interval (Fig. 2E,F). For IT neurons, the GCaMP fluorescence signal remained elevated in aiTBS-treated but not sham-treated mice for 4-8 seconds after stimulation onset, whereas activity in PT neurons dropped back to baseline during this period (Fig. 2E,F). To determine how the overall magnitude of signals arising from IT and PT neurons changed, we normalized fluorescence activity during aiTBS to the corresponding sham signals and plotted the activity across the entire 10-minute aiTBS session for each cell type (Fig. 2G). IT neurons in aiTBS- treated mice exhibited increased activity relative to sham-treated mice throughout the 10-minute aiTBS session, whereas PT neurons exhibited a trend in the opposite direction (Fig. 2H). Thus, aiTBS elicits distinct activity patterns in prefrontal IT and PT cells.

We hypothesized that the prolonged increase in IT neuron activity during an aiTBS session may potentiate activity in IT circuits to ameliorate depression-like behavior. To test this, on the day following aiTBS treatment, we used fiber photometry to record signals from dmPFC neurons during bouts of effortful activity in the tail suspension test (TST; Fig. 3A). We used this assay because the lateralized fiber optic placement was incompatible with the tight walls of the FST and other assays. Like the FST, the TST also measures effortful coping in response to an inescapable acute stressor. During the TST, IT neurons exhibited significantly higher activity at the onset of struggling epochs in aiTBS-treated compared to sham-treated mice (Fig. 3B-E). In aiTBS-treated mice, ramping IT neuron activity was evident even before the onset of struggling bouts (Fig. 3D). Moreover, there was enhanced IT activity from the onset of effortful struggling behavior to at least 15 seconds afterward (Fig. 3E). In contrast, there was no lasting effect of aiTBS on PT neuron activity associated with the onset of struggling bout (Fig. 3B). We did not observe any significant changes in activity at struggling offset in either cell type (Fig. 3C).

**Fig. 3.**
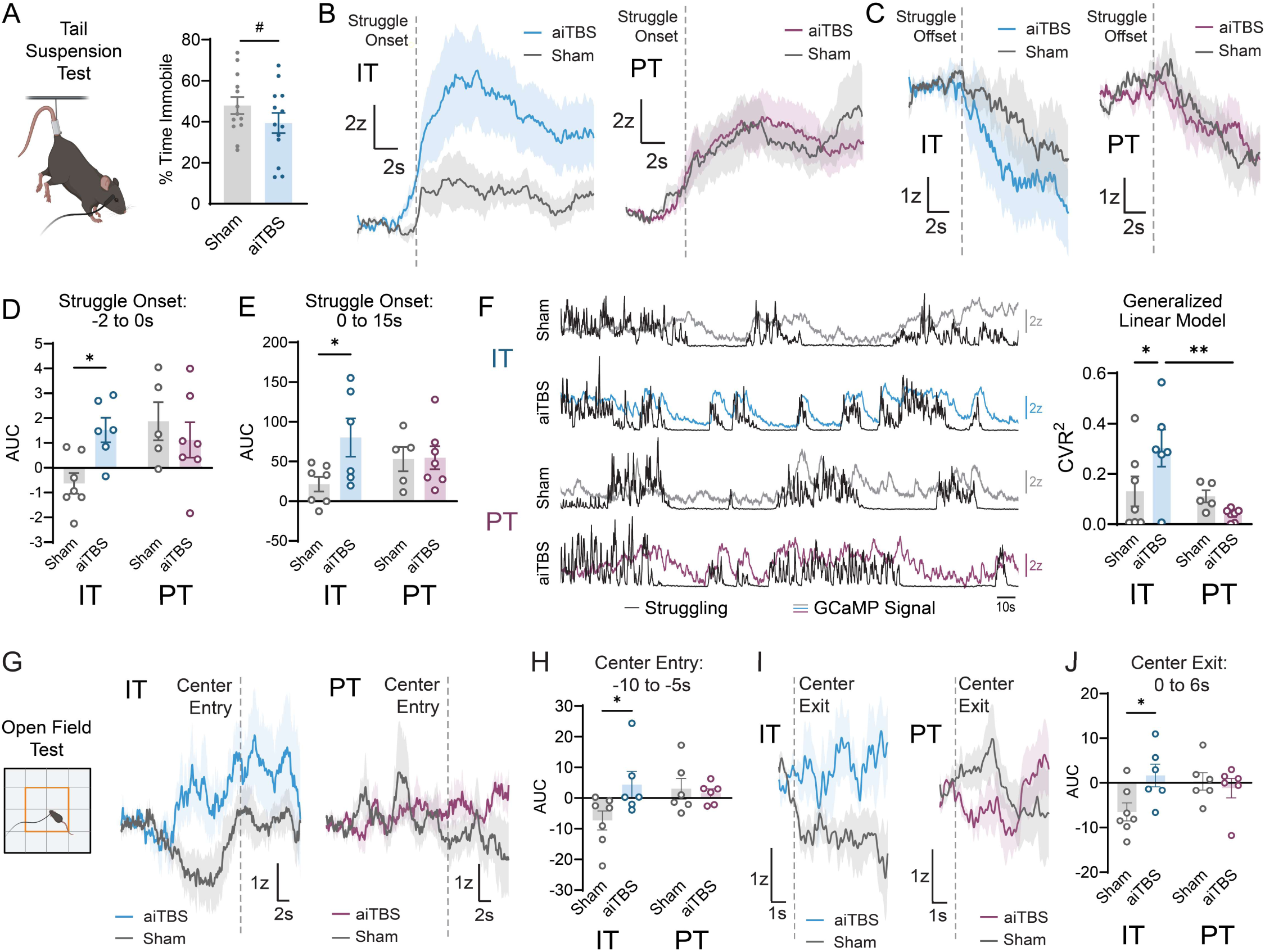
Behavioral effects of aiTBS are associated with increased IT neuron activity. (**A**) Immobility during the first 4 minutes of TST in fiber photometry mice (Sham n=13, aiTBS n=13 mice, unpaired t-test). (**B**) Fiber photometry recordings of IT and PT neurons during struggling onset in mice that received aiTBS or sham treatment. (**C**) Same as B, but recordings of neural activity during struggling offset. (**D,E**) AUC values from time windows relative to struggling onset (IT sham n=7, IT aiTBS n=6, PT sham n=5, PT aiTBS n=7 mice; two-way ANOVA with Sidak’s multiple comparisons test). (**F**) Left, representative plots showing overlay of GCaMP signal (gray or color lines) and struggling bouts during TST (black line). Right, cross-validated R^2^ values from generalized linear models fit between photometry signals and TST behavior in each animal (IT sham n=7, IT aiTBS n=6, PT sham n=5, PT aiTBS n=7 mice; two-way ANOVA with Sidak’s multiple comparisons test). (**G)** Fiber photometry recordings of Ca^2+^ fluorescence in IT and PT neurons prior to OFT center entry in aiTBS or sham-treated mice. (**H**) AUC values from -10 to -5s relative to center entry (IT sham n=7, IT aiTBS n=6, PT sham n=6, PT aiTBS n=6 mice; two-way ANOVA with Sidak’s multiple comparisons test). (**I**) Fiber photometry recordings of Ca2+ fluorescence following center exits during OFT. (**J**) AUC values from 0 to 6s following center exit (IT sham n=7, IT aiTBS n=6, PT sham n=6, PT aiTBS n=6 mice; two-way ANOVA with Sidak’s multiple comparisons test). Mouse diagrams were generated using Biorender.com. For detailed statistical analysis see Table S1. Lines and shading reflect mean ± SEM of average traces across mice. All error bars reflect mean ± SEM. #p<0.1, *p<0.05, **p<0.01.

Based on these findings, we hypothesized that aiTBS may modulate how strongly IT or PT neurons encode effortful stress coping responses. To test this, we fit a generalized linear model to examine the predictive relationship between the cell type-specific activity and an automated measure of struggling vigor for each mouse. aiTBS-treated animals exhibited an increase in the strength of the correlation between neural signals and struggling vigor for IT, but not PT neurons (Fig. 3F). Altogether, these results demonstrate that aiTBS drives plasticity in IT neurons that enhances IT neuronal activity during effortful coping behavior.

We next investigated whether the effects of aiTBS on IT and PT neuronal activity were specific to effortful coping or generalized to other behavioral responses. To test this, we recorded GCaMP fluorescence from IT and PT neurons during the OFT, which had walls wide enough to be compatible with the lateral fiber. Compared to sham-treated mice, aiTBS-treated mice had elevated IT neuron activity preceding entries to and following exits from the center of the arena (Fig. 3G-J). In contrast, aiTBS did not drive changes in PT neuron activity in either case. Consistent with our TST results, these data suggest that aiTBS alters IT, but not PT, activity-behavior relationships in a manner that may ameliorate depression- and anxiety-related behaviors.

dmPFC projection neurons, including those that fall within IT and PT classes, differentially encode aversive and rewarding stimuli and can bias approach or avoidance of these stimuli through top-down control of downstream brain regions^81^. Chronic stress can lead to pathological processing of emotional stimuli that manifests in mood and anxiety disorders^64,82^. We hypothesized that altered encoding of such stimuli may contribute to the effects of aiTBS on conflicting reward approach and threat avoidance behaviors (Fig. S4). Because the lateralized fiber implant impeded locomotion in the tight arena of the conflict assay, we instead used a head-fixed setup to present mice with cues that predicted an aversive or rewarding stimulus while recording activity from IT or PT neurons (Fig. S6). We did not observe any effect of aiTBS on IT or PT neuronal activity during either stimulus. This suggests that the aiTBS-driven behavioral effects are not a direct consequence of altered IT response to rewarding or aversive stimuli. This result further indicates that aiTBS does not universally increase IT neuron activity, but rather enhances engagement of IT neurons during specific depression-relevant behaviors.

### aiTBS reverses stress-related dendritic spine deficits in a cell type-specific manner

Converging evidence indicates that synaptic remodeling underlies both the emergence of depression and its resolution with effective treatment^83^. Atrophy of prefrontal dendritic structure, including decreased density of dendritic spines and postsynaptic proteins is a hallmark finding in depression^84^. These changes are recapitulated in rodent models of chronic stress^44,45^, and mPFC spine loss correlates with deficits in behaviors that require mPFC^85,86^. Effective antidepressant therapies can augment synapse function, promote new synapse formation and restore lost dendritic spines^49,83,87–89^. In some cases, dendritic spine elaboration is required for antidepressant behavioral responses^46^. Spine elaboration is an activity-dependent process, downstream of excitatory synaptic plasticity mechanisms proposed to mediate the effects of rTMS^9,18,19,90–95^.

We hypothesized that aiTBS-driven elaboration of dendritic spines may underlie the increased activity we observed in IT neurons during depression-related behavior. To explore this, we used a viral-genetic strategy^96^ to sparsely and brightly label the dendrites of either IT or PT neurons (Fig. 4A). Mice with cell type-specific labeling were exposed to chronic CORT and treated with aiTBS. We then imaged fluorescently labeled dmPFC IT and PT neurons using confocal microscopy (Fig. 4B,C) and quantified spine density on apical and basal dendrites, which receive synaptic input from distinct sources^97^ (Fig. 4D-G).

**Fig. 4.**
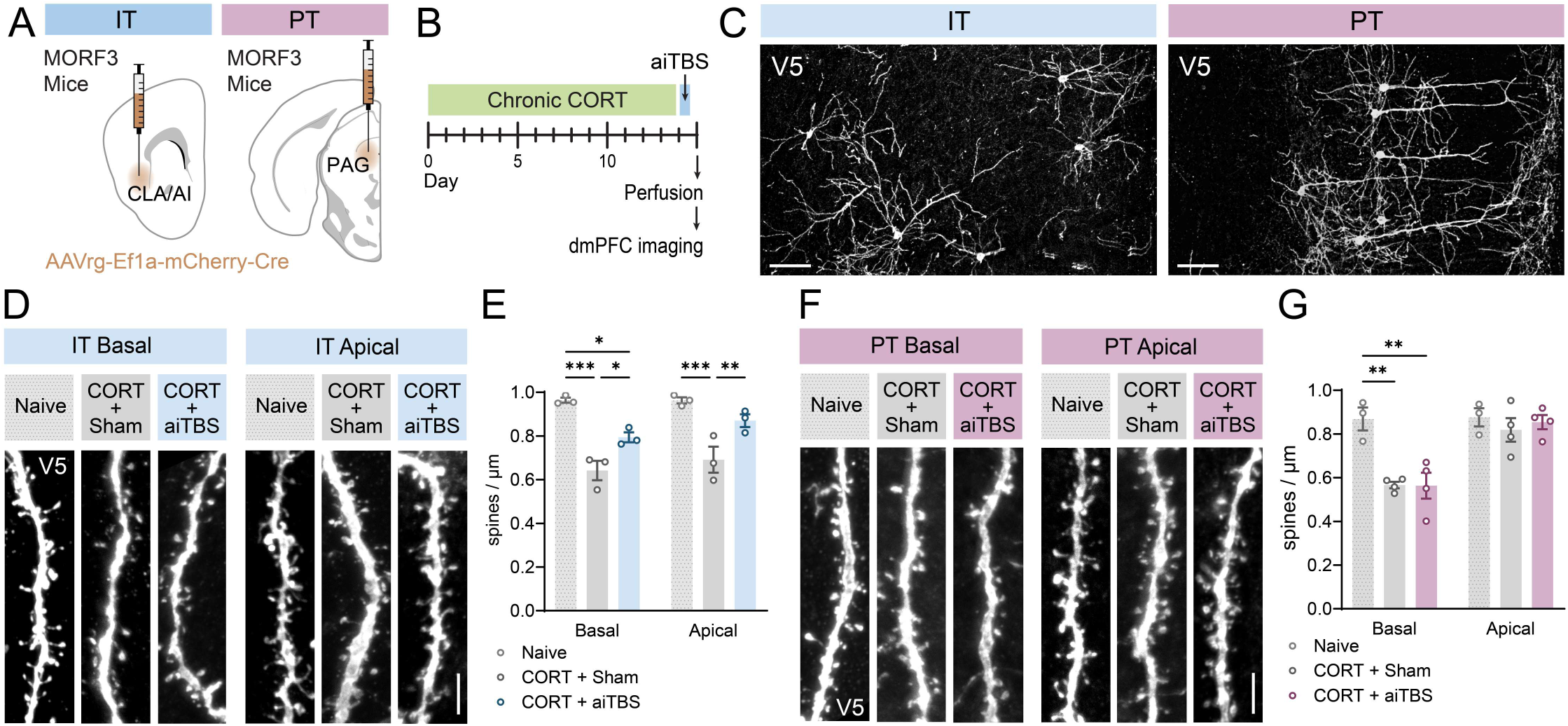
aiTBS selectively elicits dendritic spine growth in IT neurons. (**A**) Viral strategy to achieve sparse fluorescent labeling of dmPFC IT and PT neurons in MORF3 mice. (**B**) Experimental timeline. Mice were placed on CORT for 14 days, followed by a day of aiTBS treatment. The day after aiTBS treatment mice were perfused for histological analysis. (**C**) Representative images of sparsely labeled IT and PT neurons in dmPFC of MORF3 mice. Scale bars 100μm. (**D**) Representative images of dendritic spines of IT neurons in naive, CORT+sham treated, and CORT+aiTBS treated mice. Scale bar 5μm. (**E**) Average dendritic spine density on IT neurons of mice from each condition (Naive n=3, CORT+sham n=3, CORT+aiTBS n=3 mice, at least 8 dendritic segments per mouse; two-way RM ANOVA with Sidak’s multiple comparisons test). (**F**) Representative images of dendritic spines of PT neurons in naive, CORT+sham treated, and CORT+aiTBS treated mice. Scale bar 5μm. (**G**) Average dendritic spine density on PT neurons of mice from each condition (Naive n=3, CORT+sham n=4, CORT+aiTBS n=4 mice, at least 8 dendritic segments per mouse; two-way RM ANOVA with Sidak’s multiple comparisons test). For detailed statistical analysis see Table S1. All error bars reflect mean ± SEM. *p<0.05, **p<0.01, ***p<0.001.

Mice exposed to chronic CORT exhibited reduced dendritic spine density on both IT and PT neurons compared to non-stressed controls (Fig. 4D-G). In IT neurons, aiTBS treatment partially reversed loss of dendritic spines on both apical and basal dendrites (Fig. 4E). In contrast, aiTBS had no effect on PT neuron spine density (Fig. 4G), indicating that aiTBS-driven spine elaboration was specific to IT neurons. These data suggest that, like other rapid-acting antidepressant manipulations^46,49,87–89^, aiTBS may drive activity-dependent processes that promote restoration of lost dendritic spines. Our data further indicate that the effects of aiTBS are cell type-specific and could reflect strengthening or restoration of lost synaptic inputs onto IT neurons, potentially augmenting IT neuron engagement and IT circuit-mediated behavioral functions.

### Suppressing the activity of IT neurons during aiTBS blocks its antidepressant behavioral effect

Our findings suggest IT neurons are uniquely responsive to aiTBS, showing increased activity over the course of a stimulation session and enhanced activity the following day. Thus, we hypothesized that activation of these neurons during aiTBS may be necessary for subsequent changes in depression-related behavior. To test this, we used chemogenetics to selectively suppress IT neuron activity during aiTBS. We first expressed the inhibitory DREADD (designer receptor exclusively activated by designer drugs) hM4Di^98^ specifically in dmPFC IT neurons (Fig. 5A, S7). We then injected mice with the hM4Di ligand clozapine-N-oxide (CNO) to suppress the activity of these neurons during aiTBS. To evaluate the effectiveness of this strategy, we injected a subset of mice with CNO prior to a single 10-minute aiTBS session. Afterwards, we examined expression of the immediate early gene Fos, a marker of recent neuronal activity (Fig. 5B). In mice injected with CNO, we observed an ∼75% reduction in Fos expression in hM4Di-expressing IT neurons compared to mCherry-only controls (Fig. 5B). This indicates that our chemogenetic manipulation effectively suppressed aiTBS-driven activity in IT neurons.

**Fig. 5.**
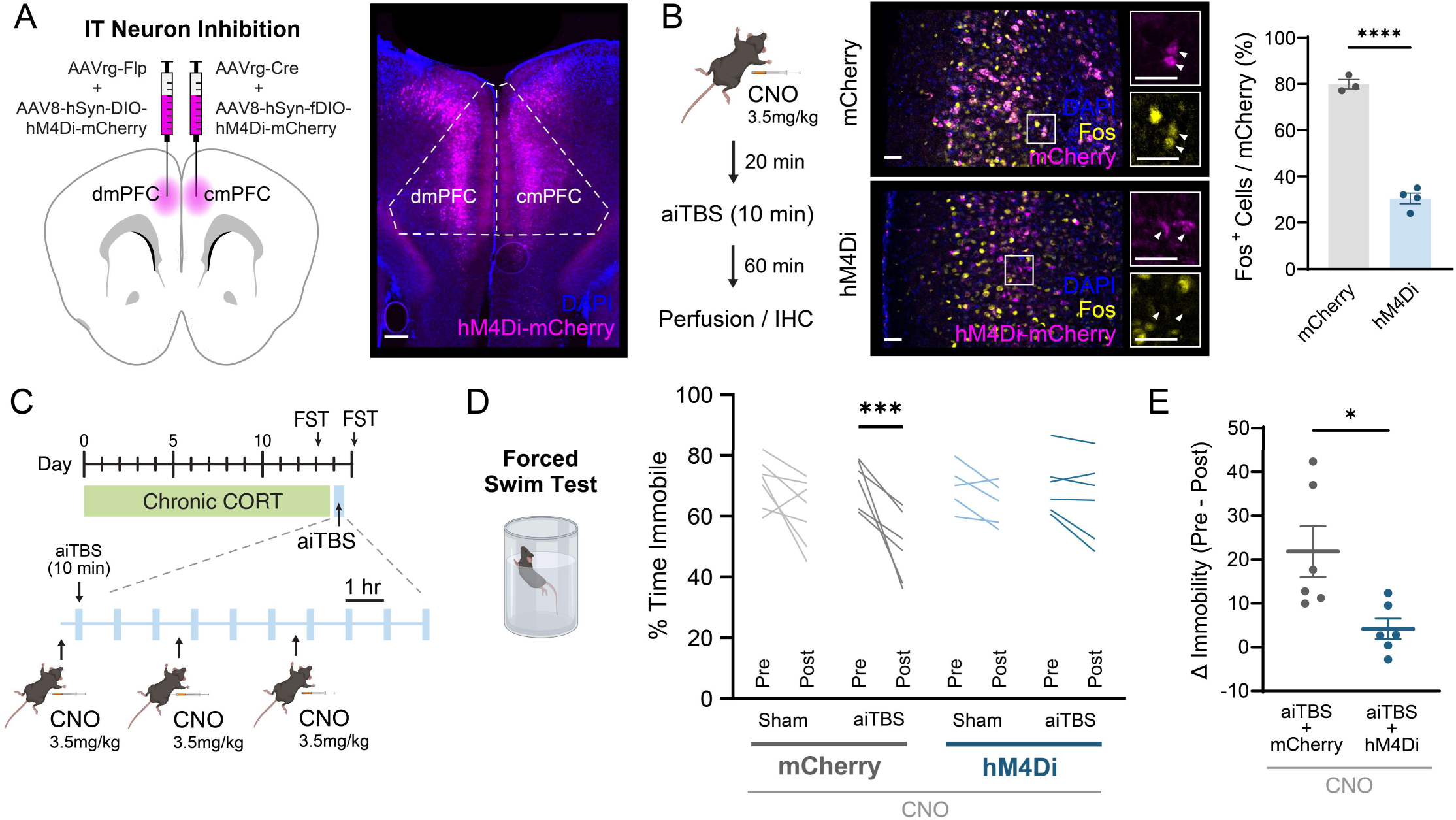
Inhibition of IT neurons blocks antidepressant effects of aiTBS. (**A**) Left, viral strategy for cell-type specific chemogenetic inhibition of IT neurons in dmPFC. Right, representative image of hM4Di expression in dmPFC IT neurons. Scale bar 200μm. (**B**) Fos expression in dmPFC following neuronal inhibition with hM4Di. Middle, representative images of Fos and mCherry colocalization. Scale bars 40μm. Right, percent of mCherry^+^ neurons expressing Fos in mice expressing hM4Di (mCherry n=3, hM4Di n=4 mice; unpaired t-test). (**C**) Experimental strategy for inhibition of IT neurons during aiTBS. Following viral injections, mice were placed on CORT for 14 days and treated with aiTBS. Mice were injected with CNO prior to the first, fourth, and seventh aiTBS session of the day. FST was performed the day before and the day after aiTBS treatment. (**D**) Effects of IT neuron inhibition on immobility in the forced swim test (Sham+mCherry n=7, aiTBS+mCherry n=6, Sham+hM4Di n=5, aiTBS+hM4Di n=6 mice; three-way RM ANOVA with Sidak’s multiple comparisons test). (**E**) Effects of IT neuron inhibition on aiTBS-induced change in immobility (aiTBS+mCherry n=6, aiTBS+hM4Di n=6 mice, unpaired t-test). Mouse diagrams were generated using Biorender.com. For detailed statistical analysis see Table S1. All error bars reflect mean ± SEM. *p<0.05, ***p<0.001, ****p<0.0001.

Next, to test whether IT activation is required to generate the antidepressant effects of aiTBS, we chemogenetically suppressed activity in IT neurons throughout the entire course of aiTBS treatment (Fig. 5C). In mCherry control animals, aiTBS but not sham treatment significantly reduced immobility during the FST compared to the pre-TMS baseline (Fig. 5D). This aligns with our previous behavioral effect in the FST and confirms that the effect is not eliminated by CNO itself. In contrast, in mice expressing hM4Di in IT neurons, aiTBS did not affect immobility relative to the pre-TMS baseline (Fig. 5D). In line with this, mice expressing hM4Di exhibited a smaller change in immobility from pre to post aiTBS treatment compared to mCherry controls (Fig. 5E). Thus, suppressing the activity of IT neurons during aiTBS treatment is sufficient to prevent antidepressant-like changes in effortful coping behavior. Together our findings demonstrate that clinical rTMS protocols such as aiTBS drive cell type-specific modifications in IT projection neurons and aiTBS-driven activation of this projection class is required to reverse a behavioral deficit associated with chronic stress.

## Discussion

Non-invasive brain stimulation promises to treat brain disorders by resolving disturbances in circuit function, but how specific circuits or cell types are affected is not known. Our study establishes a novel rodent model of rTMS and sheds light on the cell type-resolved antidepressant mechanism of aiTBS. We discovered that aiTBS, the first rapid rTMS protocol for depression, rescues behavioral deficits induced by chronic stress via a cell type-specific neural mechanism. Compared to PT cells, IT cells are uniquely sensitive to aiTBS, showing a gradual increase in activity across a session. A single day of aiTBS restores lost dendritic spines and enhances neural activity during effortful behavior in prefrontal IT, but not PT, neurons. Neural activity in local prefrontal IT neurons is required during rTMS to produce the antidepressant-like behavioral effects of aiTBS.

This study represents a significant advance in our understanding of how rTMS can modulate specific cortical cell-types to produce clinically useful effects. Prior studies revealed that rTMS modifies gene expression and produces structural and functional changes reminiscent of synaptic long-term potentiation (LTP) or depression (LTD)^18,19,99–101^. In some cases these effects were specific to excitatory or inhibitory neurons^18–22,102,103^. While important, these early studies were based on models that lacked clinical face validity because they used non-clinical protocols, anesthetized animals, were not focal and suprathreshold, or investigated non-clinical targets. Our work overcomes these limitations and links cell-type specific neural activation during rTMS to subsequent behavioral effects. IT and PT projection classes are a highly conserved circuit architecture across cortical regions and between species^73,74^. We find that IT neurons are uniquely responsive to aiTBS-driven plasticity. Their unique response properties may arise as a function of their specific local and long range connectivity, molecular expression profiles, or synaptic properties^72,75–79,104^. The extent to which each of these features contribute remains to be explored. Our findings are highly consistent with a companion manuscript co-submitted with this report, which provides additional evidence that aiTBS selectively modulates IT circuits^105^.

There is growing interest in understanding the circuit changes that produce rapid resolution of depression^46,49,106,107^. Depression is characterized by heterogeneous disturbances in cognitive and emotional brain networks that can arise at the intersection of genetic and environmental factors, including chronic stress^13,43,108,109^. It is important to note that rodent behavioral assays cannot directly prove relevance to symptoms of depression, but we used assays that evoke cognitive and emotional processes disrupted in chronic stress. The PFC is a critical hub in these networks and is highly sensitive to chronic stress. Hypofunction of prefrontal projection neurons and the resulting aberrant activity patterns are a point of convergence for factors that promote or alleviate depressed states^46,67,69^.

Our findings suggest IT neurons are a novel target for the development of precision medicine interventions for depression and other conditions, by selective modulation using aiTBS or other brain stimulation strategies. It is possible that different stimulation protocols, or combination of rTMS with pharmacologic agents, will drive distinct effects, promoting plasticity in other cell types. In the clinic, combining rTMS with modulation of dopaminergic and noradrenergic activity via psychostimulants may enhance antidepressant outcomes^110,111^. This suggests combination therapy may promote plasticity in circuits that may not otherwise respond to rTMS. Beyond depression, aiTBS and other rTMS protocols are being studied in the treatment of other neurological and psychiatric disorders, including addiction, schizophrenia, dementia, and chronic pain ^112^. Our work sets the foundation for the discovery of noninvasive cell-type specific treatment protocols that are targeted to dysfunctional circuit elements of these unique symptoms and disorders.

## Acknowledgements

We thank Conor Liston, Shane Johnson, Joshua Barry, Carlos Cepeda, Michael Levine, Liqun Luo and Baljit Khakh for helpful discussions while designing and planning experiments and helpful comments on the manuscript. We also thank the UCLA Eli and Edythe Broad Stem Cell Research Center (BSCRC) microscopy core for access to imaging facilities.

## Author contributions

Conceptualization: MWG, AFL, LAD, SAW

Methodology: MWG, ASE, AQ, AQK, GAW, YY, HL, AFL, LAD, SAW Investigation: MWG, AQ, ASE, SARM, CBK, MS, OHW, AH, JPR, HL, LAD, SAW

Visualization: MWG

Funding acquisition: MWG, AFL, LAD, SAW Resources: AFL, LAD, SAW

Project administration: MWG, LAD, SAW Supervision: MWG, AFL, LAD, SAW Writing – original draft: MWG

Writing – review & editing: MWG, HL, AFL, LAD, SAW

## Data and materials availability

All data are available in the main text or the supplementary materials.

## Supplementary Materials

Document S1: Figs. S1 to S7 and Table S1

## STAR Methods

### EXPERIMENTAL MODEL

Male and female C57BL/6 (Jackson Laboratories #000664 or #005304) or MORF3 mice (Jackson Laboratories #035403) at least 8 weeks of age were used for all experiments. Mice were maintained on a 12-hour light cycle (lights on 7am-7pm) in a temperature- and humidity-controlled animal facility. All animal procedures were approved by the University of California, Los Angeles Chancellor’s Animal Research Committee.

### METHOD DETAILS

#### Surgery

Mice were anesthetized with 3% isoflurane until loss of righting reflex and transferred to a stereotaxic surgical setup, where anesthesia was maintained with 1-2% isoflurane. The scalp was cleaned with three alternating swabs of betadine and 70% ethanol. 2% Lidocaine was injected under the scalp as a local anesthetic. A small incision was made in the scalp. For viral injections, a small hole was drilled above the injection target and a hamilton syringe loaded with virus was lowered to the correct stereotaxic coordinate. Virus was pressure injected at 100nL/minute, and following completion of the injection the syringe was left in place for 7 minutes and then slowly removed from the skull. Coordinates used for injections were as follows, in mm relative to bregma: dmPFC: AP 1.45, ML ±0.35, DV -1.85; PAG: AP -4.0, ML -0.45, DV -3.0; CLA/AI: AP 1.3, ML -3.25, DV -4.2.

While animals were on the stereotaxic rig, we used stereotaxic coordinates to mark the location for targeting of the TMS coil hotspot. For motor response experiments, a small marker dot was placed above the approximate hindlimb region of the motor cortex using stereotaxic coordinates (AP -1.0, ML +1.5) to use for alignment of the magnet. For dmPFC stimulation and behavioral experiments in Figures 1, 5, and S2-4, the dot was placed at AP 1.45mm, ML 0mm (on the midline) to promote bilateral stimulation. For unilateral fiber photometry and dendritic spine experiments in Figures 2-4, and S6, the dot was placed at AP 1.45mm, ML -0.35mm to stimulate above the tip of the fiber optic cannula.

All mice were also implanted with skull bars to allow head fixation during sham or TMS treatment. To do so, muscles on the posterior region of the skull were carefully removed with scissors and a scalpel blade to reveal the skull surface. Light scoring of the skull was performed with a drill, and the area was cleaned with ethanol. A 3D-printed plastic skull bar was then positioned posterior to the skull and cemented in place with metabond (Patterson Dental Company, 5533559, 5533492, S371). Care was taken to avoid buildup of metabond near the site of stimulation. Mice were pre- and then post-operatively treated with subcutaneous injections of carprofen (50mg/kg) daily for three days following surgery.

For fiber photometry experiments, skin and muscle tissue behind the eye was carefully removed on the side of the skull directly lateral to the target location. A drill was used to clear away bone on the side of the skull, and a 4mm long 400nm optic fiber was slowly inserted through this hole from the lateral direction until it reached the coordinate for dmPFC. Tissue surrounding the fiber insertion site was dried using compressed air. The fiber was then cemented in place with metabond followed by installation of a skull bar as above.

#### Chronic Stress

##### Chronic corticosterone administration

Corticosterone (Sigma) was dissolved in 100% ethanol at 10mg/mL, and this was diluted with drinking water from the animal facility to achieve a final concentration of 0.1mg/mL. Mice received this solution in their home cages in place of normal water for 14 days. Control animals received normal drinking water, consistent with previous studies^46^.

##### Unpredictable Chronic Mild Stress

Mice were exposed to two stressors daily for 21 days. Stressors included restraint stress (1-2hr), removal of bedding (2-4hr), ¼ inch of lukewarm water in place of bedding (1-2hr), wet bedding (1-2hr), tilted cage (2-4hr), rapid air flow in cage (10min), light cycle disturbance (overnight), and food/water restriction (overnight). Stressors were randomized, and the same stressor was never presented on two consecutive days^55^.

#### Behavioral Experiments

##### Open Field Test & Elevated Zero Maze

Mice were gently placed in the corner of an open field arena (50x50cm) facing the wall of the arena. Mice were allowed to explore the arena for 10 minutes while their behavior was recorded with an overhead camera. Mice were then removed from the arena, briefly placed in their home cage, and then placed at the entry to the closed arm of the EZM. Mice were allowed to explore the EZM for 10 minutes while their behavior was recorded with an overhead camera. Mice were then returned to their home cage. The OFT and EZM were cleaned with 70% ethanol between each mouse. Time spent in specific zones for each assay was quantified using Biobserve Viewer software.

##### Forced swim test

2-liter glass beakers were filled with 25-26°C water and placed against a white background in a dimly lit room. Mice were gently lowered into the water and their behavior recorded during a 6- minute window from the side of the beaker. Mice were then removed from the beakers, dried, and placed back in their home cage. Water was changed between each animal. Immobility during the last 4 minutes of the assay was quantified using DBscorer^113^, which classifies struggling behavior by calculating the change in the area of the frame taken up by the mouse across time. To ensure consistent baseline levels of immobility in mice undergoing FST both before and after aiTBS (Figures S1 and 5), only mice with more than 56.5% immobility on the pretest were used for subsequent aiTBS or sham treatment. This was equivalent to 1 standard deviation below the mean immobility of all stress mice in Figure 1E.

##### Sucrose preference test

Mice were placed in individual chambers (25cm L x 10cm W x 12.5 cm H) with two glass drinking bottles (Bio-Serv #9019/9015) suspended on one short end of the chamber^61^. Mice were water deprived for 20 hours before each habituation and test session. Mice were first habituated to the setup for 4 hours with both bottles filled with normal drinking water. Mice were then placed back in the chamber for another 4 hour habituation session, this time with 1% sucrose in one of the bottles. The following day the testing session began. Mice were given free choice between normal drinking water and 1% sucrose in water for 4 hours. The position of the bottles were switched halfway through to eliminate effects of a potential side preference. Bottles were weighed before and after the 4 hour session, and sucrose preference was calculated as (grams sucrose solution consumed) / (total liquid consumed) x 100. 3 test sessions were run before CORT administration, and the average sucrose preference across these sessions was considered the baseline sucrose preference. For these animals, CORT administration was extended to 18 days to account for the potential loss of stress effects once the CORT was removed for water deprivation. Following CORT, mice were water deprived for one day and another habituation session was performed with 1% sucrose in one of the bottles, followed by a test session one day later. Mice were treated with aiTBS or sham the following day. One day after aiTBS or sham treatment, mice were given another habituation session with 1% sucrose in one of the bottles, followed by a final test session the next day.

##### Sinking platform test

An abbreviated version of the sinking platform test^60^ was performed to allow the rapid assessment of persistence behavior shortly after the administration of CORT and subsequent aiTBS treatment. A 32-gallon plastic container 22” in diameter (Brute H-1045) was filled with approximately 10 inches of 32°C water. A custom-built platform 3.35”x4.8” was positioned level with the surface of the water in the locations described in Figure S3D. Locations were varied around the tank, with some located in the center of the tank. On the first day, mice were first exposed to four escape trials, in which climbing the platform resulted in removal from the tank. The same day, a progressive ratio was introduced consisting of an increasing number of failure trials prior to the final escape trial. A failure trial consisted of sinking the platform and repositioning it after the mouse climbed it, requiring the mouse to swim to the new location. The second day, two additional training sessions were performed with 4 and 7 failure trials before the escape trial, respectively. Latency per platform during training was calculated as the total time swimming prior to climbing the final platform divided by the number of platforms climbed. After these sessions, the same day, mice were subjected to a test session consisting of exclusively failure trials. The total number of platforms climbed within 5 minutes was reported for each mouse. Mice were treated with TMS the following day, and the day 2 protocol was repeated, with two training sessions followed by a final test session.

##### Approach-avoidance conflict assay

Mice were food deprived for 4 days prior to the beginning of habituation, receiving free food access for 1 hour per day. Behavior was performed in an operant chamber with a shock floor (Lafayette Instruments). One quarter of the floor was covered by a plexiglass platform, and in the opposite corner from this platform was a reward pump. The reward pump, fitted with an IR sensor, was set to deliver rewards following head entry at a randomized variable interval schedule averaging 30 seconds per interval, regardless of tone presentation, as previously described^63^. Mice were habituated to the setup and reward delivery during the final two days of CORT administration to allow for the immediate start of training following aiTBS or sham treatment. The day after treatment, mice received 3 baseline tones with no shock followed by 9 tones which co- terminated with a 2-second mild foot shock (0.13mA). Tones were 4kHz, 70dB, and 30 seconds in length and were separated by randomized intervals between 80 and 150 seconds. Reward delivery continued on the variable interval schedule regardless of tone presentation. On the second day of training, mice received 9 additional tone-shock pairings. On the retrieval day, three tones were presented without foot shocks. Videos of behavior were obtained using a Point Gray Chameleon3 camera (Teledyne FLIR). Point-tracking was performed in DeepLabCut^114^, and behavior was analyzed using BehaviorDEPOT^115^. Retrieval videos for two mice (one sham, one aiTBS) were irreversibly corrupted due to technical difficulties and were not able to be analyzed.

#### Transcranial Magnetic Stimulation

The rodent TMS coil was assembled as previously described and validated^34–36^. It was connected to a MagStim Rapid^2^ Plus1 stimulator, which was used to power the coil and deliver clinical stimulation protocols. To prevent overheating of the magnet during chronic, high-frequency stimulation, sheets of copper were placed directly against the four long sides of the coil housing, and ¼ inch copper tubing was wrapped around the magnet as shown in Figure 1A. This setup was designed to optimize heat transfer between the liquid in the copper tubing and the epoxy- coated coil. An aquarium pump was submerged in a solution of 30% CaCl2 in water maintained at -40°C and was used to pump this solution through the copper tubing beginning at the start of the stimulation session. This maintained the surface of the magnet at approximately room temperature for the duration of a stimulation session.

For motor response measurements, awake mice were scruffed in a way that largely immobilized their limbs, and the hotspot of the magnet was positioned over the marked coordinate. Fine adjustments in head position were made until reliable left hindlimb movement was obtained with single pulses. The magnet intensity was then adjusted to find the minimum threshold at which 50% of pulses resulted in motor responses, which is how motor threshold is assessed in humans. Across five mice, we estimated the motor threshold to be between 50 and 55% of the maximum stimulator output.

For chronic treatment, mice were head-fixed on a running wheel by clamping surgical hemostats onto their skull bars and gently placing these hemostats into a custom 3D-printed hemostat holder. Mice were habituated to head fixation for at least two days prior to treatment. Prior to each stimulation session, the stereotaxic marker dot indicating the position of dmPFC was aligned to a dot indicating the focal ‘hotspot’ of the TMS coil using a fine 3-axis manipulator. The coil was then lowered such that the epoxy coating at the ‘hotspot’ of the magnet pressed directly against the mouse’s skull.

The TMS coil produces an audible clicking noise and slight vibrations during stimulation. To ensure sham-treated animals had an identical experience to experimental animals, with the absence of magnetic stimulation, sham mice were always run simultaneously on an identical head fixation setup directly adjacent to the setup for TMS-treated animals (Fig. S1). This ensured nearly identical auditory stimuli between groups. In addition, the lid of a 50mL conical tube, which has a texture similar to the epoxy coating of the real magnet, was pressed directly against the head of the sham mouse. To account for mechanical sensations during stimulation, the heavy cable connecting the TMS coil to the stimulator was draped over the sham coil to produce identically- timed vibrations which were transmitted through the sham coil.

Accelerated iTBS was performed using clinically identical stimulation parameters as previously described^23,24,41^. Ten 50Hz bursts of 3 pulses, separated by 200ms intervals, were delivered within a two second window. We refer to this as a ‘train’. Trains were delivered every 10 seconds for a total of 60 trains across a 10-minute stimulation session. 10 sessions were delivered throughout the treatment day, with a session occuring once per hour. The stimulation intensity was maintained at 42% of the maximum stimulator output, in line with clinical iTBS, which is typically delivered at a power between 80 and 90% of the motor threshold. Mice were returned to their home cage between stimulation sessions.

#### Fiber Photometry

To record neural activity in IT and PT neurons, 200nL of AAVrg-Ef1a-mCherry-IRES-Cre (Addgene #55632, 2.2x10^13^ vg/mL) was injected in either right dmPFC or left PAG, respectively. 400nL of AAV1-Syn-FLEX-jGCaMP7f (Addgene #104492, 2.1x10^13^ vg/mL) was then injected into left dmPFC. Viruses were injected at least 4 weeks prior to the start of recordings. Fiber photometry signals were recorded using a Tucker-Davis Technologies (TDT) RZ10x processor in combination with the TDT Synapse software. The 465nm channel representing calcium-driven changes in GCaMP fluorescence was recorded simultaneously with a 405nm isosbestic channel, which does not fluctuate with calcium activity. Prior to recording, the light output of each channel for each mouse was adjusted to a signal of approximately 80mV.

##### Recording during rTMS

To control for auditory-driven neural activity not directly linked to magnetic stimulation, the sound of the magnet was recorded and played constantly through computer speakers for the duration of combined TMS and fiber photometry recording. Recordings during iTBS treatment in Figure 2E were limited to the first session of the day to prevent the constant auditory noise from affecting behavioral outcomes the following day. A TTL signal connecting the stimulator to the fiber photometry system allowed alignment of the stimulation and the photometry signal.

##### Pavlovian stimuli

Beginning two days prior to TMS treatment, fiber photometry mice were head-fixed and underwent pavlovian condition for aversive and rewarding stimuli. Aversive stimuli consisted of a 5-second 7kHz tone at approximately 75dB co-terminating with a 0.5-second air puff. The air puff spout was positioned approximately 7mm from the eye. Rewarding stimuli consisted of a 5-second 2.9kHz tone at approximately 75dB co-terminating with a 0.5-second delivery of 25% sweetened condensed milk via a peristaltic pump. The reward spout was positioned directly in the mouse’s mouth to ensure reward delivery occurred at the exact same time for each stimulus and to guarantee the mouse received a reward on each trial. On the first day of conditioning, mice received 6 rewarding stimuli followed by 6 additional rewarding stimuli randomly interspersed with 6 aversive stimuli. On the second day of conditioning, mice received 6 stimuli of each type, randomly interspersed. Stimuli were separated by a randomized interval between 30 and 60 seconds each. The day after TMS treatment, mice received 6 stimuli of each type, randomly interspersed, while photometry signals were recorded. Tones, air puffs and rewards were aligned with fiber photometry signals via TTL pulses delivered to the fiber photometry system.

##### Tail suspension test and Open Field Test

Because FST was not compatible with the lateral fiber optic implant without risking animal safety, we instead used the tail suspension test to record neural activity during struggling behavior. A small plastic sheath was placed on the mouse’s tail to prevent tail climbing, and the end of the tail was secured with a piece of tape. The other end of this piece of tape was wrapped around a wooden dowel. Behavior was recorded for 6.5 minutes. TTL pulses were delivered every 30 seconds to the photometry system via the video recording computer to allow alignment between behavior and photometry signals. The same TTL setup was used to align OFT behavior with photometry.

#### Histology & Immunohistochemistry

Following completion of experiments, mice were anesthetized with isoflurane and transcardially perfused with 15mL of 0.1M phosphate-buffered saline (PBS, Gibco Sciences) followed by 15mL of 4% paraformaldehyde (PFA, Sigma) in PBS. Brains were dissected from the skull and post- fixed in PFA for 24 hours, after which they were transferred to either PBS + 0.01% sodium azide (for vibratome sectioning) or 30% sucrose in PBS (for cryosectioning). Once brains for cryosectioning no longer floated in 30% sucrose, brains were embedded in Optimum Cutting Temperature (OCT; Tissue Tek) and frozen at -80°C, followed by sectioning on a cryostat at 60μm. Other brains were sectioned on a vibratome at 60μm in PBS. Sections were stored in cryoprotectant at -20°C until staining. For validation of fiber tracts and viral expression, sections were DAPI stained, mounted, and imaged with a 5x objective on a Leica DM6 B scanning microscope.

##### MORF3 Dendritic Spine Imaging

Heterozygous MORF3 mice were injected with 200nL AAVrg-Ef1a-mCherry-IRES-Cre (Addgene #55632, 2.2x10^13^ vg/mL) in either claustrum/agranular insula (CLA/AI) or PAG to label dmPFC IT and PT neurons, respectively. CLA/AI was used as an IT target instead of contralateral PFC to promote sparser labeling and limit background from dense back-projecting axons. Mice were also implanted with a skull bar to allow aiTBS or sham treatment. After at least one week of recovery from surgery, mice were placed on CORT for 14 days, followed by one day of aiTBS or sham treatment. The following day, mice were perfused. Vibratome sections of these brains were collected at 60μm.

To stain for V5, the epitope tag used to label Cre-expressing cells in MORF3 mice, sections were washed 3 x 10 minutes in PBS, followed by 1 hour in blocking solution containing 3% Normal Donkey Serum (NDS), 3% Bovine Serum Albumin (BSA), and 0.5% Triton-X100 in PBS. Sections were then transferred to blocking solution containing primary antibody (Rabbit anti-V5, Bethyl A190-120A, 1:500) for 3 nights, shaking at 4°C. Sections were washed 3 x 10 minutes in PBS and transferred to blocking solution containing secondary antibody (Donkey anti-rabbit, Alexa 647 conjugate, Jackson ImmunoResearch 711-605-152, 1:5000) for one night, shaking at 4°C. Sections were washed 3 x 10 minutes in PBS, stained with DAPI (1:4000), and mounted.

Spine images were collected using a 63x oil immersion objective on a Zeiss LSM700 confocal microscope. A z-stack extending above and below the planes in which the dendrite was visible was obtained to ensure the capture of all dendritic spines. The interval between z-steps was 0.45μm. At least 8 dendrites were collected per brain, split between apical and basal dendrites.

#### Chemogenetic Inhibition of IT Neurons

To achieve bilateral inhibition of dmPFC cells in each hemisphere projecting to the contralateral hemisphere, a four-virus intersectional strategy was used. In left dmPFC, mice were injected with a mixture of 400nL AAV8-hSyn-DIO-hM4D(Gi)-mCherry (Addgene #44362, 2x10^13^ vg/mL) and 200nL AAVrg-Ef1a-FlpO (Addgene #55637, 1.6x10^13^ vg/mL). In right dmPFC, mice were injected with a mixture of 400nL AAV8-hSyn-fDIO-hM4D(Gi)-mCherry (Addgene #154867, 2.5x10^13^ vg/mL) and 200nL AAVrg-Ef1a-Cre (Addgene #55636, 2.2x10^13^ vg/mL). Consistent with previous studies of contralaterally-projecting mPFC neurons^71,104^, this approach yielded viral expression that was largely restricted to layers 2/3 and 5a of dmPFC, with some expression in deeper layers (Fig. 5A, S7). Control animals received AAV8-hSyn-DIO-mCherry (Addgene #50459, 2.4x10^13^ vg/mL) and AAV8-Ef1a-fDIO-mCherry (Addgene #114471, 2.3x10^13^ vg/mL) in place of Hm4Di viruses. All mice were implanted with a skull bar to allow aiTBS or sham treatment. 4 weeks of viral expression were allowed prior to treatment.

To assess the effects of hM4Di on neuronal activity during aiTBS, mice expressing either hM4Di or mCherry were injected with CNO dihydrochloride (3.5mg/kg in 0.9% saline, Fisher Scientific 63-295-0). 20 minutes later, mice underwent one session of aiTBS treatment. 60 minutes following treatment, mice were perfused for histological analysis of neuronal activation via Fos. 60μm cryosections were washed 3 x 10 minutes in PBS, followed by 1 hour in blocking solution containing 10% Normal Donkey Serum (NDS) and 0.3% Triton-X100 in PBS. Sections were transferred to blocking solution containing primary antibody (Rabbit anti-Fos, Synaptic Systems 226008, 1:1000) for 3 nights, shaking at 4°C. Sections were washed 3 x 10 minutes in PBS and transferred to blocking solution containing secondary antibody (Donkey anti-rabbit, Alexa 647 conjugate, Jackson ImmunoResearch 711-605-152, 1:1000) for 2 hours at room temperature. Sections were washed 3 x 10 minutes in PBS, stained with DAPI (1:4000), and mounted. Z stacks with an interval of 1μm were collected using a 10x objective on a Leica STELLARIS confocal microscope. Colocalization between Fos and mCherry was analyzed in ImageJ^116^.

For behavioral chemogenetics experiments, mice were placed on CORT for 14 days. On the 14th day, mice underwent FST to obtain a pre-TMS baseline. The next day, mice underwent iTBS or sham treatment. All mice were injected with 3.5mg/kg CNO 20 minutes prior to the first, fourth, and seventh sessions of iTBS. The following day, mice underwent FST as above.

### QUANTIFICATION AND STATISTICAL ANALYSIS

Statistical tests were performed in GraphPad Prism. Details of statistical testing and results are available in the figure legends and Table S1. For all experiments n represents an individual mouse. All error bars represent mean±SEM. Shading on photometry plots represents mean±SEM calculated using the average trace from all animals in each condition. In all figures ^#^p<0.1, *p<0.05, **p<0.01, ***p<0.001, ****p<0.0001. Data distributions were tested in Prism using D’Agostino & Pearson, Anderson-Darling, Shapiro-Wilk, and Kolmogorov-Smirnov tests prior to application of tests assuming a normal distribution. Welch’s t-test was used in place of an unpaired t-test when the F test revealed a significant difference in variance between conditions.

#### Fiber Photometry Analysis

Fiber photometry analysis was performed in MATLAB based on example code provided by TDT as previously described^71^. In brief, raw 1kHz recordings were downsampled by a factor of 10. The MATLAB polyfit function was used to fit a curve between the 405 and 465 signals for each recording. The predicted 465 values based on the 405 channel were subtracted from the raw 465 signal to produce the signal used for analysis. The mean and standard deviation of a period prior to each event of interest was used to baseline the signal for each event, and the z-score based on this baseline period was calculated across each event. For recordings during TMS, the baseline period was -2 to 0 seconds relative to the start of the stimulation train. For pavlovian stimuli, the baseline period was -10 to 0 seconds relative to the start of the tone, as we have previously described for auditory stimuli^71^.

For OFT and TST behaviors not time-locked to a particular stimulus, events were only analyzed if the end of the previous event occurred prior to the baseline of the next period (i.e. if the mouse had just stopped struggling less than 5 seconds before the onset of a new struggling event, this onset event was not analyzed because the previous event would confound the baseline activity).

Baseline periods were chosen to maximize the number of included events while maintaining the ability to capture potential changes in activity prior to the start of the event. For TST, the baseline period was -5 to -2 seconds relative to the onset or offset of struggling. Baseline for OFT center entry was -20 to -15 seconds relative to entry, consistent with our previous studies of innate avoidance behavior^71^. Baseline for OFT center exits was -1 to 0 seconds, as entries to the center often lasted only 1 to 3 seconds. All behavioral epochs were annotated by an expert observer blinded to the corresponding fiber photometry signal.

All statistical analyses in Figures 2, 3, and S6 were performed by calculating the average trace for all events from each animal. Each animal’s trace was separately used to calculate the area under the curve for the time interval of interest, and statistical comparisons were performed across animals. To visualize these signals, traces from each animal were smoothed with a moving average of 0.2 seconds, and the mean ± SEM of all animals’ average traces was plotted.

For generalized linear modeling of TST behavior and photometry signals in Figure 3F, the z-score of each animal’s photometry signal was calculated across the entire session. DBscorer was used to generate a plot of the change in area from each frame taken up by the mouse, which correlates with the vigor at which each mouse was struggling at any point in time. A generalized linear model with a poisson distribution was fit for each animal using 10-fold cross validation, with the photometry signal as the input variable and the behavior as the response variable. The cross- validated R^2^ value from this model was compared across animals.

#### Dendritic Spine Analysis

Spines were counted in ImageJ^116^ by an expert observer blinded to condition. The simple neurite tracer plugin^117^ was used to trace the length of the dendrite in 3D space, and the spines along the traced dendrite were counted manually by scrolling through the z stack to ensure no spines were missed. Values are reported as the number of spines counted divided by the length of the traced dendrite. The average value from apical and basal dendrites from each brain was used for statistical comparisons between conditions.

## References

1. Rajasethupathy, P., Ferenczi, E., and Deisseroth, K. (2016). Targeting Neural Circuits. Cell 165, 524–534. 10.1016/j.cell.2016.03.047.

2. Barker, A.T., Jalinous, R., and Freeston, I.L. (1985). Non-invasive magnetic stimulation of human motor cortex. Lancet Lond. Engl. 1, 1106–1107. 10.1016/s0140-6736(85)92413-4.

3. Rothwell, J.C., Thompson, P.D., Day, B.L., Boyd, S., and Marsden, C.D. (1991). Stimulation of the human motor cortex through the scalp. Exp. Physiol. 76, 159–200. 10.1113/expphysiol.1991.sp003485.

4. Lefaucheur, J.-P. (2009). Methods of therapeutic cortical stimulation. Neurophysiol. Clin. Clin. Neurophysiol. 39, 1–14. 10.1016/j.neucli.2008.11.001.

5. 5. Lefaucheur, J.-P., Aleman, A., Baeken, C., Benninger, D.H., Brunelin, J., Di Lazzaro, V., Filipović, S.R., Grefkes, C., Hasan, A., Hummel, F.C., et al. (2020). Evidence-based guidelines on the therapeutic use of repetitive transcranial magnetic stimulation (rTMS): An update (2014-2018). Clin. Neurophysiol. Off. J. Int. Fed. Clin. Neurophysiol. 131, 474–528. 10.1016/j.clinph.2019.11.002.

6. George, M.S., Wassermann, E.M., Williams, W.A., Callahan, A., Ketter, T.A., Basser, P., Hallett, M., and Post, R.M. (1995). Daily repetitive transcranial magnetic stimulation (rTMS) improves mood in depression. Neuroreport 6, 1853–1856. 10.1097/00001756-199510020-00008.

7. Pascual-Leone, A., Rubio, B., Pallardó, F., and Catalá, M.D. (1996). Rapid-rate transcranial magnetic stimulation of left dorsolateral prefrontal cortex in drug-resistant depression. Lancet Lond. Engl. 348, 233–237. 10.1016/s0140-6736(96)01219-6.

8. Acevedo, N., Bosanac, P., Pikoos, T., Rossell, S., and Castle, D. (2021). Therapeutic Neurostimulation in Obsessive-Compulsive and Related Disorders: A Systematic Review. Brain Sci. 11, 948. 10.3390/brainsci11070948.

9. Pell, G.S., Roth, Y., and Zangen, A. (2011). Modulation of cortical excitability induced by repetitive transcranial magnetic stimulation: Influence of timing and geometrical parameters and underlying mechanisms. Prog. Neurobiol. 93, 59–98. 10.1016/j.pneurobio.2010.10.003.

10. 10. Suppa, A., Asci, F., and Guerra, A. (2022). Transcranial magnetic stimulation as a tool to induce and explore plasticity in humans. In Handbook of Clinical Neurology, A. Quartarone, M. F. Ghilardi, and F. Boller, eds. (Elsevier), pp. 73–89. 10.1016/B978-0-12-819410-2.00005-9.

11. George, M.S., Lisanby, S.H., Avery, D., McDonald, W.M., Durkalski, V., Pavlicova, M., Anderson, B., Nahas, Z., Bulow, P., Zarkowski, P., et al. (2010). Daily Left Prefrontal Transcranial Magnetic Stimulation Therapy for Major Depressive Disorder: A Sham- Controlled Randomized Trial. Arch. Gen. Psychiatry 67, 507–516. 10.1001/archgenpsychiatry.2010.46.

12. Mirman, A.M., Corlier, J., Wilson, A.C., Tadayonnejad, R., Marder, K.G., Pleman, C.M., Krantz, D.E., Wilke, S.A., Levitt, J.G., Ginder, N.D., et al. (2022). Absence of early mood improvement as a robust predictor of rTMS nonresponse in major depressive disorder. Depress. Anxiety 39, 123–133. 10.1002/da.23237.

13. George, M.S., Ketter, T.A., and Post, R.M. (1994). Prefrontal cortex dysfunction in clinical depression. Depression 2, 59–72. 10.1002/depr.3050020202.

14. Fitzgerald, P.B., Laird, A.R., Maller, J., and Daskalakis, Z.J. (2008). A meta-analytic study of changes in brain activation in depression. Hum. Brain Mapp. 29, 683–695. 10.1002/hbm.20426.

15. Elbau, I.G., Lynch, C.J., Downar, J., Vila-Rodriguez, F., Power, J.D., Solomonov, N., Daskalakis, Z.J., Blumberger, D.M., and Liston, C. (2023). Functional Connectivity Mapping for rTMS Target Selection in Depression. Am. J. Psychiatry 180, 230–240. 10.1176/appi.ajp.20220306.

16. Young, I.M., Taylor, H.M., Nicholas, P.J., Mackenzie, A., Tanglay, O., Dadario, N.B., Osipowicz, K., Davis, E., Doyen, S., Teo, C., et al. (2023). An agile, data-driven approach for target selection in rTMS therapy for anxiety symptoms: Proof of concept and preliminary data for two novel targets. Brain Behav. 13, e2914. 10.1002/brb3.2914.

17. Brown, K.E., Neva, J.L., Ledwell, N.M., and Boyd, L.A. (2014). Use of transcranial magnetic stimulation in the treatment of selected movement disorders. Degener. Neurol. Neuromuscul. Dis. 4, 133–151. 10.2147/DNND.S70079.

18. Vlachos, A., Müller-Dahlhaus, F., Rosskopp, J., Lenz, M., Ziemann, U., and Deller, T. (2012). Repetitive magnetic stimulation induces functional and structural plasticity of excitatory postsynapses in mouse organotypic hippocampal slice cultures. J. Neurosci. Off. J. Soc. Neurosci. 32, 17514–17523. 10.1523/JNEUROSCI.0409-12.2012.

19. Lenz, M., Platschek, S., Priesemann, V., Becker, D., Willems, L.M., Ziemann, U., Deller, T., Müller-Dahlhaus, F., Jedlicka, P., and Vlachos, A. (2015). Repetitive magnetic stimulation induces plasticity of excitatory postsynapses on proximal dendrites of cultured mouse CA1 pyramidal neurons. Brain Struct. Funct. 220, 3323–3337. 10.1007/s00429-014-0859-9.

20. Funke, K., and Benali, A. (2011). Modulation of cortical inhibition by rTMS - findings obtained from animal models. J. Physiol. 589, 4423–4435. 10.1113/jphysiol.2011.206573.

21. Lenz, M., Galanis, C., Müller-Dahlhaus, F., Opitz, A., Wierenga, C.J., Szabó, G., Ziemann, U., Deller, T., Funke, K., and Vlachos, A. (2016). Repetitive magnetic stimulation induces plasticity of inhibitory synapses. Nat. Commun. 7, 10020. 10.1038/ncomms10020.

22. Trippe, J., Mix, A., Aydin-Abidin, S., Funke, K., and Benali, A. (2009). θ burst and conventional low-frequency rTMS differentially affect GABAergic neurotransmission in the rat cortex. Exp. Brain Res. 199, 411–421. 10.1007/s00221-009-1961-8.

23. Cole, E.J., Stimpson, K.H., Bentzley, B.S., Gulser, M., Cherian, K., Tischler, C., Nejad, R., Pankow, H., Choi, E., Aaron, H., et al. (2020). Stanford Accelerated Intelligent Neuromodulation Therapy for Treatment-Resistant Depression. Am. J. Psychiatry 177, 716–726. 10.1176/appi.ajp.2019.19070720.

24. Cole, E.J., Phillips, A.L., Bentzley, B.S., Stimpson, K.H., Nejad, R., Barmak, F., Veerapal, C., Khan, N., Cherian, K., Felber, E., et al. (2022). Stanford Neuromodulation Therapy (SNT): A Double-Blind Randomized Controlled Trial. Am. J. Psychiatry 179, 132–141. 10.1176/appi.ajp.2021.20101429.

25. 25. van Rooij, S.J.H., Arulpragasam, A.R., McDonald, W.M., and Philip, N.S. (2024). Accelerated TMS - moving quickly into the future of depression treatment. Neuropsychopharmacology 49, 128–137. 10.1038/s41386-023-01599-z.

26. Cohen, D., and Cuffin, B.N. (1991). Developing a More Focal Magnetic Stimulator. Part I: Some Basic Principles. J. Clin. Neurophysiol. 8, 102.

27. Wilson, M.T., Tang, A.D., Iyer, K., McKee, H., Waas, J., and Rodger, J. (2018). The challenges of producing effective small coils for transcranial magnetic stimulation of mice. Biomed. Phys. Eng. Express 4, 037002. 10.1088/2057-1976/aab525.

28. Gersner, R., Kravetz, E., Feil, J., Pell, G., and Zangen, A. (2011). Long-Term Effects of Repetitive Transcranial Magnetic Stimulation on Markers for Neuroplasticity: Differential Outcomes in Anesthetized and Awake Animals. J. Neurosci. 31, 7521–7526. 10.1523/JNEUROSCI.6751-10.2011.

29. Tang, A.D., Lowe, A.S., Garrett, A.R., Woodward, R., Bennett, W., Canty, A.J., Garry, M.I., Hinder, M.R., Summers, J.J., Gersner, R., et al. (2016). Construction and Evaluation of Rodent-Specific rTMS Coils. Front. Neural Circuits 10. 10.3389/fncir.2016.00047.

30. Vahabzadeh-Hagh, A.M., Muller, P.A., Gersner, R., Zangen, A., and Rotenberg, A. (2012). Translational Neuromodulation: Approximating Human Transcranial Magnetic Stimulation Protocols in Rats. Neuromodulation Technol. Neural Interface 15, 296–305. 10.1111/j.1525-1403.2012.00482.x.

31. Rotenberg, A., Muller, P.A., Vahabzadeh-Hagh, A.M., Navarro, X., López-Vales, R., Pascual-Leone, A., and Jensen, F. (2010). Lateralization of forelimb motor evoked potentials by transcranial magnetic stimulation in rats. Clin. Neurophysiol. 121, 104–108. 10.1016/j.clinph.2009.09.008.

32. Alekseichuk, I., Mantell, K., Shirinpour, S., and Opitz, A. (2019). Comparative modeling of transcranial magnetic and electric stimulation in mouse, monkey, and human. NeuroImage 194, 136–148. 10.1016/j.neuroimage.2019.03.044.

33. Parthoens, J., Verhaeghe, J., Servaes, S., Miranda, A., Stroobants, S., and Staelens, S. (2016). Performance Characterization of an Actively Cooled Repetitive Transcranial Magnetic Stimulation Coil for the Rat. Neuromodulation Technol. Neural Interface 19, 459–468. 10.1111/ner.12387.

34. Meng, Q., Jing, L., Badjo, J.P., Du, X., Hong, E., Yang, Y., Lu, H., and Choa, F.-S. (2018). A novel transcranial magnetic stimulator for focal stimulation of rodent brain. Brain Stimulat. 11, 663–665. 10.1016/j.brs.2018.02.018.

35. Meng, Q., Nguyen, H., Vrana, A., Baldwin, S., Li, C.Q., Giles, A., Wang, J., Yang, Y., and Lu, H. (2022). A high-density theta burst paradigm enhances the aftereffects of transcranial magnetic stimulation: Evidence from focal stimulation of rat motor cortex. Brain Stimulat. 15, 833–842. 10.1016/j.brs.2022.05.017.

36. Cermak, S., Meng, Q., Peng, K., Baldwin, S., Mejías-Aponte, C.A., Yang, Y., and Lu, H. (2020). Focal transcranial magnetic stimulation in awake rats: enhanced glucose uptake in deep cortical layers. J. Neurosci. Methods 339, 108709. 10.1016/j.jneumeth.2020.108709.

37. Bagherzadeh, H., Meng, Q., Deng, Z.-D., Lu, H., Hong, E., Yang, Y., and Choa, F.-S. (2022). Angle-tuned coils: attractive building blocks for TMS with improved depth-spread performance. J. Neural Eng. 19, 10.1088/1741-2552/ac697c.

38. Tennant, K.A., Adkins, D.L., Donlan, N.A., Asay, A.L., Thomas, N., Kleim, J.A., and Jones, T.A. (2011). The Organization of the Forelimb Representation of the C57BL/6 Mouse Motor Cortex as Defined by Intracortical Microstimulation and Cytoarchitecture. Cereb. Cortex 21, 865–876. 10.1093/cercor/bhq159.

39. Li, C.-X., and Waters, R.S. (1991). Organization of the Mouse Motor Cortex Studied by Retrograde Tracing and Intracortical Microstimulation (ICMS) Mapping. Can. J. Neurol. Sci. 18, 28–38. 10.1017/S0317167100031267.

40. Huang, Y.-Z., Edwards, M.J., Rounis, E., Bhatia, K.P., and Rothwell, J.C. (2005). Theta burst stimulation of the human motor cortex. Neuron 45, 201–206. 10.1016/j.neuron.2004.12.033.

41. Raj, K.S., Geoly, A.D., Veerapal, C., Gholmieh, M., Toosi, P., Espil, F.M., Batail, J.-M., Kratter, I.H., and Williams, N.R. (2024). Pilot study of stanford neuromodulation therapy (SNT) for bipolar depression. Brain Stimul. Basic Transl. Clin. Res. Neuromodulation 17, 321–323. 10.1016/j.brs.2024.03.002.

42. Suzuki, M., Furihata, R., Konno, C., Kaneita, Y., Ohida, T., and Uchiyama, M. (2018). Stressful events and coping strategies associated with symptoms of depression: A Japanese general population survey. J. Affect. Disord. 238, 482–488. 10.1016/j.jad.2018.06.024.

43. Tafet, G.E., and Nemeroff, C.B. (2016). The Links Between Stress and Depression: Psychoneuroendocrinological, Genetic, and Environmental Interactions. J. Neuropsychiatry Clin. Neurosci. 28, 77–88. 10.1176/appi.neuropsych.15030053.

44. Radley, J.J., Rocher, A.B., Miller, M., Janssen, W.G.M., Liston, C., Hof, P.R., McEwen, B.S., and Morrison, J.H. (2006). Repeated stress induces dendritic spine loss in the rat medial prefrontal cortex. Cereb. Cortex N. Y. N 1991 *16*, 313–320. 10.1093/cercor/bhi104.

45. Radley, J.J., Rocher, A.B., Rodriguez, A., Ehlenberger, D.B., Dammann, M., McEwen, B.S., Morrison, J.H., Wearne, S.L., and Hof, P.R. (2008). REPEATED STRESS ALTERS DENDRITIC SPINE MORPHOLOGY IN THE RAT MEDIAL PREFRONTAL CORTEX. J. Comp. Neurol. 507, 1141–1150. 10.1002/cne.21588.

46. Moda-Sava, R.N., Murdock, M.H., Parekh, P.K., Fetcho, R.N., Huang, B.S., Huynh, T.N., Witztum, J., Shaver, D.C., Rosenthal, D.L., Alway, E.J., et al. (2019). Sustained rescue of prefrontal circuit dysfunction by antidepressant-induced spine formation. Science 364, eaat8078. 10.1126/science.aat8078.

47. Fetcho, R.N., Parekh, P.K., Chou, J., Kenwood, M., Chalençon, L., Estrin, D.J., Johnson, M., and Liston, C. (2024). A stress-sensitive frontostriatal circuit supporting effortful reward-seeking behavior. Neuron 112, 473–487.e4. 10.1016/j.neuron.2023.10.020.

48. Orzechowska, A., Zajączkowska, M., Talarowska, M., and Gałecki, P. (2013). Depression and ways of coping with stress: a preliminary study. Med. Sci. Monit. Int. Med. J. Exp. Clin. Res. 19, 1050–1056. 10.12659/MSM.889778.

49. Li, N., Lee, B., Liu, R.-J., Banasr, M., Dwyer, J.M., Iwata, M., Li, X.-Y., Aghajanian, G., and Duman, R.S. (2010). mTOR-dependent synapse formation underlies the rapid antidepressant effects of NMDA antagonists. Science 329, 959–964. 10.1126/science.1190287.

50. Kwan, A.C., Olson, D.E., Preller, K.H., and Roth, B.L. (2022). The neural basis of psychedelic action. Nat. Neurosci. 25, 1407–1419. 10.1038/s41593-022-01177-4.

51. Gourley, S.L., and Taylor, J.R. (2009). Recapitulation and Reversal of a Persistent Depression-like Syndrome in Rodents. Curr. Protoc. Neurosci. 49, 9.32.1-9.32.11. 10.1002/0471142301.ns0932s49.

52. Marks, W., Fournier, N.M., and Kalynchuk, L.E. (2009). Repeated exposure to corticosterone increases depression-like behavior in two different versions of the forced swim test without altering nonspecific locomotor activity or muscle strength. Physiol. Behav. 98, 67–72. 10.1016/j.physbeh.2009.04.014.

53. Wróbel, A., Serefko, A., Poleszak, E., and Rechberger, T. (2016). Fourteen-day administration of corticosterone may induce detrusor overactivity symptoms. Int. Urogynecology J. 27, 1713–1721. 10.1007/s00192-016-3027-3.

54. Nollet, M. (2021). Models of Depression: Unpredictable Chronic Mild Stress in Mice. Curr. Protoc. 1, e208. 10.1002/cpz1.208.

55. Rawat, R., Tunc-Ozcan, E., McGuire, T.L., Peng, C.-Y., and Kessler, J.A. (2022). Ketamine activates adult-born immature granule neurons to rapidly alleviate depression-like behaviors in mice. Nat. Commun. 13, 2650. 10.1038/s41467-022-30386-5.

56. Gorman, J.M. (1996). Comorbid depression and anxiety spectrum disorders. Depress. Anxiety 4, 160–168. 10.1002/(SICI)1520-6394(1996)4:4<160::AID-DA2>3.0.CO;2-J.

57. Tuinstra, D., Percifield, C., Stilwell, K., Plattner, A., Edwards, E., Sanders, W., and Koval, M. (2022). Treatment of anxiety symptoms in patients receiving rTMS for treatment resistant depression. Psychiatry Res. Commun. 2, 100014. 10.1016/j.psycom.2021.100014.

58. Stanton, C.H., Holmes, A.J., Chang, S.W.C., and Joormann, J. (2019). From Stress to Anhedonia: Molecular Processes through Functional Circuits. Trends Neurosci. 42, 23–42. 10.1016/j.tins.2018.09.008.

59. Prevot, T.D., Misquitta, K.A., Fee, C., Newton, D.F., Chatterjee, D., Nikolova, Y.S., Sibille, E., and Banasr, M. (2019). Residual avoidance: A new, consistent and repeatable readout of chronic stress-induced conflict anxiety reversible by antidepressant treatment. Neuropharmacology 153, 98–110. 10.1016/j.neuropharm.2019.05.005.

60. Floris, G., Godar, S.C., Braccagni, G., Piras, I.S., Ravens, A., Zanda, M.T., Huentelman, M.J., and Bortolato, M. (2024). The sinking platform test: a novel paradigm to measure persistence in animal models. Neuropsychopharmacology 49, 1373–1382. 10.1038/s41386-024-01827-0.

61. Liu, M.-Y., Yin, C.-Y., Zhu, L.-J., Zhu, X.-H., Xu, C., Luo, C.-X., Chen, H., Zhu, D.-Y., and Zhou, Q.-G. (2018). Sucrose preference test for measurement of stress-induced anhedonia in mice. Nat. Protoc. 13, 1686–1698. 10.1038/s41596-018-0011-z.

62. Letkiewicz, A.M., Kottler, H.C., Shankman, S.A., and Cochran, A.L. (2023). Quantifying aberrant approach-avoidance conflict in psychopathology: A review of computational approaches. Neurosci. Biobehav. Rev. 147, 105103. 10.1016/j.neubiorev.2023.105103.

63. Zeidler, Z., Gómez, M.F., Gupta, T.A., Shari, M., Wilke, S.A., and DeNardo, L.A. (2024). Prefrontal dopamine circuits are required for active avoidance learning but not for fear learning. Preprint at bioRxiv, 10.1101/2024.05.02.592069.

64. Russo, S.J., and Nestler, E.J. (2013). The brain reward circuitry in mood disorders. Nat. Rev. Neurosci. 14, 609–625. 10.1038/nrn3381.

65. Warden, M.R., Selimbeyoglu, A., Mirzabekov, J.J., Lo, M., Thompson, K.R., Kim, S.-Y., Adhikari, A., Tye, K.M., Frank, L.M., and Deisseroth, K. (2012). A prefrontal cortex– brainstem neuronal projection that controls response to behavioural challenge. Nature 492, 428–432. 10.1038/nature11617.

66. Hare, B.D., Shinohara, R., Liu, R.J., Pothula, S., DiLeone, R.J., and Duman, R.S. (2019). Optogenetic stimulation of medial prefrontal cortex Drd1 neurons produces rapid and long-lasting antidepressant effects. Nat. Commun. 10, 223. 10.1038/s41467-018-08168-9.

67. Shrestha, P., Mousa, A., and Heintz, N. (2015). Layer 2/3 pyramidal cells in the medial prefrontal cortex moderate stress induced depressive behaviors. eLife 4, e08752. 10.7554/eLife.08752.

68. Cai, Y., Ge, J., and Pan, Z.Z. (2024). The projection from dorsal medial prefrontal cortex to basolateral amygdala promotes behaviors of negative emotion in rats. Front. Neurosci. 18. 10.3389/fnins.2024.1331864.

69. Johnson, S.B., Lingg, R.T., Skog, T.D., Hinz, D.C., Romig-Martin, S.A., Viau, V., Narayanan, N.S., and Radley, J.J. (2022). Activity in a prefrontal-periaqueductal gray circuit overcomes behavioral and endocrine features of the passive coping stress response. Proc. Natl. Acad. Sci. U. S. A. 119, e2210783119. 10.1073/pnas.2210783119.

70. Wilke, S.A., Lavi, K., Byeon, S., Donohue, K.C., and Sohal, V.S. (2022). Convergence of Clinically Relevant Manipulations on Dopamine-Regulated Prefrontal Activity Underlying Stress Coping Responses. Biol. Psychiatry 91, 810–820. 10.1016/j.biopsych.2021.11.008.

71. Gongwer, M.W., Klune, C.B., Couto, J., Jin, B., Enos, A.S., Chen, R., Friedmann, D., and DeNardo, L.A. (2023). Brain-wide projections and differential encoding of prefrontal neuronal classes underlying learned and innate threat avoidance. J. Neurosci. 10.1523/JNEUROSCI.0697-23.2023.

72. Gao, L., Liu, S., Gou, L., Hu, Y., Liu, Y., Deng, L., Ma, D., Wang, H., Yang, Q., Chen, Z., et al. (2022). Single-neuron projectome of mouse prefrontal cortex. Nat. Neurosci. 25, 515–529. 10.1038/s41593-022-01041-5.

73. Gerfen, C.R., Economo, M.N., and Chandrashekar, J. (2018). Long distance projections of cortical pyramidal neurons. J. Neurosci. Res. 96, 1467–1475. 10.1002/jnr.23978.

74. Harris, K.D., and Shepherd, G.M.G. (2015). The neocortical circuit: themes and variations. Nat. Neurosci. 18, 170–181. 10.1038/nn.3917.

75. Anastasiades, P.G., and Carter, A.G. (2021). Circuit organization of the rodent medial prefrontal cortex. Trends Neurosci. 44, 550–563. 10.1016/j.tins.2021.03.006.

76. Anastasiades, P.G., Collins, D.P., and Carter, A.G. (2021). Mediodorsal and Ventromedial Thalamus Engage Distinct L1 Circuits in the Prefrontal Cortex. Neuron 109, 314–330.e4. 10.1016/j.neuron.2020.10.031.

77. Anastasiades, P.G., Marlin, J.J., and Carter, A.G. (2018). Cell-Type Specificity of Callosally Evoked Excitation and Feedforward Inhibition in the Prefrontal Cortex. Cell Rep. 22, 679– 692. 10.1016/j.celrep.2017.12.073.

78. Collins, D.P., Anastasiades, P.G., Marlin, J.J., and Carter, A.G. (2018). Reciprocal Circuits Linking the Prefrontal Cortex with Dorsal and Ventral Thalamic Nuclei. Neuron 98, 366–379.e4. 10.1016/j.neuron.2018.03.024.

79. 79. Yao, Z., van Velthoven, C.T.J., Kunst, M., Zhang, M., McMillen, D., Lee, C., Jung, W., Goldy, J., Abdelhak, A., Aitken, M., et al. (2023). A high-resolution transcriptomic and spatial atlas of cell types in the whole mouse brain. Nature 624, 317–332. 10.1038/s41586-023-06812-z.

80. Shepherd, G.M.G. (2013). Corticostriatal connectivity and its role in disease. Nat. Rev. Neurosci. 14, 278–291. 10.1038/nrn3469.

81. Pastor, V., and Medina, J.H. (2021). Medial prefrontal cortical control of reward- and aversion-based behavioral output: Bottom-up modulation. Eur. J. Neurosci. 53, 3039– 3062. 10.1111/ejn.15168.

82. 82. Coley, A.A., Padilla-Coreano, N., Patel, R., and Tye, K.M. (2021). Chapter Seven - Valence processing in the PFC: Reconciling circuit-level and systems-level views. In International Review of Neurobiology What does Medial Frontal Cortex Signal During Behavior? Insights from Behavioral Neurophysiology., A. T. Brockett, L. M. Amarante, M. Laubach, and M. R. Roesch, eds. (Academic Press), pp. 171–212. 10.1016/bs.irn.2020.12.002.

83. Parekh, P.K., Johnson, S.B., and Liston, C. (2022). Synaptic mechanisms regulating mood state transitions in depression. Annu. Rev. Neurosci. 45, 581–601. 10.1146/annurev-neuro-110920-040422.

84. Kang, H.J., Voleti, B., Hajszan, T., Rajkowska, G., Stockmeier, C.A., Licznerski, P., Lepack, A., Majik, M.S., Jeong, L.S., Banasr, M., et al. (2012). Decreased expression of synapse-related genes and loss of synapses in major depressive disorder. Nat. Med. 18, 1413–1417. 10.1038/nm.2886.

85. Dias-Ferreira, E., Sousa, J.C., Melo, I., Morgado, P., Mesquita, A.R., Cerqueira, J.J., Costa, R.M., and Sousa, N. (2009). Chronic stress causes frontostriatal reorganization and affects decision-making. Science 325, 621–625. 10.1126/science.1171203.

86. Liston, C., Miller, M.M., Goldwater, D.S., Radley, J.J., Rocher, A.B., Hof, P.R., Morrison, J.H., and McEwen, B.S. (2006). Stress-induced alterations in prefrontal cortical dendritic morphology predict selective impairments in perceptual attentional set-shifting. J. Neurosci. Off. J. Soc. Neurosci. 26, 7870–7874. 10.1523/JNEUROSCI.1184-06.2006.

87. Bessa, J.M., Ferreira, D., Melo, I., Marques, F., Cerqueira, J.J., Palha, J.A., Almeida, O.F.X., and Sousa, N. (2009). The mood-improving actions of antidepressants do not depend on neurogenesis but are associated with neuronal remodeling. Mol. Psychiatry 14, 764–773, 739. 10.1038/mp.2008.119.

88. Hesselgrave, N., Troppoli, T.A., Wulff, A.B., Cole, A.B., and Thompson, S.M. (2021). Harnessing psilocybin: antidepressant-like behavioral and synaptic actions of psilocybin are independent of 5-HT2R activation in mice. Proc. Natl. Acad. Sci. U. S. A. 118, e2022489118. 10.1073/pnas.2022489118.

89. Shao, L.-X., Liao, C., Gregg, I., Davoudian, P.A., Savalia, N.K., Delagarza, K., and Kwan, A.C. (2021). Psilocybin induces rapid and persistent growth of dendritic spines in frontal cortex in vivo. Neuron 109, 2535–2544.e4. 10.1016/j.neuron.2021.06.008.

90. Huang, Y.-Z., Chen, R.-S., Rothwell, J.C., and Wen, H.-Y. (2007). The after-effect of human theta burst stimulation is NMDA receptor dependent. Clin. Neurophysiol. Off. J. Int. Fed. Clin. Neurophysiol. 118, 1028–1032. 10.1016/j.clinph.2007.01.021.

91. Teo, J.T.H., Swayne, O.B., and Rothwell, J.C. (2007). Further evidence for NMDA- dependence of the after-effects of human theta burst stimulation. Clin. Neurophysiol. Off. J. Int. Fed. Clin. Neurophysiol. 118, 1649–1651. 10.1016/j.clinph.2007.04.010.

92. Selby, B., MacMaster, F.P., Kirton, A., and McGirr, A. (2019). d-cycloserine blunts motor cortex facilitation after intermittent theta burst transcranial magnetic stimulation: A double-blind randomized placebo-controlled crossover study. Brain Stimulat. 12, 1063–1065. 10.1016/j.brs.2019.03.026.

93. Brown, J.C., Yuan, S., DeVries, W.H., Armstrong, N.M., Korte, J.E., Sahlem, G.L., Carpenter, L.L., and George, M.S. (2021). NMDA-Receptor Agonist Reveals LTP-like Properties of 10-Hz rTMS in the Human Motor Cortex. Brain Stimulat. 14, 619–621. 10.1016/j.brs.2021.03.016.

94. Thickbroom, G.W. (2007). Transcranial magnetic stimulation and synaptic plasticity: experimental framework and human models. Exp. Brain Res. 180, 583–593. 10.1007/s00221-007-0991-3.

95. Ridding, M.C., and Ziemann, U. (2010). Determinants of the induction of cortical plasticity by non-invasive brain stimulation in healthy subjects. J. Physiol. 588, 2291–2304. 10.1113/jphysiol.2010.190314.

96. Veldman, M.B., Park, C.S., Eyermann, C.M., Zhang, J.Y., Zuniga-Sanchez, E., Hirano, A.A., Daigle, T.L., Foster, N.N., Zhu, M., Langfelder, P., et al. (2020). Brainwide Genetic Sparse Cell Labeling to Illuminate the Morphology of Neurons and Glia with Cre-Dependent MORF Mice. Neuron 108, 111–127.e6. 10.1016/j.neuron.2020.07.019.

97. Little, J.P., and Carter, A.G. (2012). Subcellular synaptic connectivity of layer 2 pyramidal neurons in the medial prefrontal cortex. J. Neurosci. Off. J. Soc. Neurosci. 32, 12808– 12819. 10.1523/JNEUROSCI.1616-12.2012.

98. Armbruster, B.N., Li, X., Pausch, M.H., Herlitze, S., and Roth, B.L. (2007). Evolving the lock to fit the key to create a family of G protein-coupled receptors potently activated by an inert ligand. Proc. Natl. Acad. Sci. 104, 5163–5168. 10.1073/pnas.0700293104.

99. Amer, A., and Martin, J.H. (2022). Repeated motor cortex theta-burst stimulation produces persistent strengthening of corticospinal motor output and durable spinal cord structural changes in the rat. Brain Stimulat. 15, 1013–1022. 10.1016/j.brs.2022.07.005.

100. Lee, C.-W., Chu, M.-C., Wu, H.-F., Chung, Y.-J., Hsieh, T.-H., Chang, C.-Y., Lin, Y.-C., Lu, T.-Y., Chang, C.-H., Chi, H., et al. (2023). Different synaptic mechanisms of intermittent and continuous theta-burst stimulations in a severe foot-shock induced and treatment-resistant depression in a rat model. Exp. Neurol. 362, 114338. 10.1016/j.expneurol.2023.114338.

101. Tokay, T., Holl, N., Kirschstein, T., Zschorlich, V., and Köhling, R. (2009). High-frequency magnetic stimulation induces long-term potentiation in rat hippocampal slices. Neurosci. Lett. 461, 150–154. 10.1016/j.neulet.2009.06.032.

102. Benali, A., Trippe, J., Weiler, E., Mix, A., Petrasch-Parwez, E., Girzalsky, W., Eysel, U.T., Erdmann, R., and Funke, K. (2011). Theta-Burst Transcranial Magnetic Stimulation Alters Cortical Inhibition. J. Neurosci. 31, 1193–1203. 10.1523/JNEUROSCI.1379-10.2011.

103. Murphy, S.C., Palmer, L.M., Nyffeler, T., Müri, R.M., and Larkum, M.E. (2016). Transcranial magnetic stimulation (TMS) inhibits cortical dendrites. eLife 5, e13598. 10.7554/eLife.13598.

104. Anastasiades, P.G., Boada, C., and Carter, A.G. (2019). Cell-Type-Specific D1 Dopamine Receptor Modulation of Projection Neurons and Interneurons in the Prefrontal Cortex. Cereb. Cortex 29, 3224–3242. 10.1093/cercor/bhy299.

105. Johnson, S.B., Rocks, D., Chalencon, L., Akgul, G., Elbau, I., Estrin, D., Johnson, K., Zhang, R., Lenz, A., Mikofsky, R., et al. Fronto-insular circuit mechanisms of accelerated intermittent theta burst stimulation. Co-submitted with this report.

106. Chen, T., Cheng, L., Ma, J., Yuan, J., Pi, C., Xiong, L., Chen, J., Liu, H., Tang, J., Zhong, Y., et al. (2023). Molecular mechanisms of rapid-acting antidepressants: New perspectives for developing antidepressants. Pharmacol. Res. 194, 106837. 10.1016/j.phrs.2023.106837.

107. Krystal, J.H., Kavalali, E.T., and Monteggia, L.M. (2024). Ketamine and rapid antidepressant action: new treatments and novel synaptic signaling mechanisms. Neuropsychopharmacology 49, 41–50. 10.1038/s41386-023-01629-w.

108. Koenigs, M., and Grafman, J. (2009). The functional neuroanatomy of depression: distinct roles for ventromedial and dorsolateral prefrontal cortex. Behav. Brain Res. 201, 239–243. 10.1016/j.bbr.2009.03.004.

109. Zhao, M.-Z., Song, X.-S., and Ma, J.-S. (2021). Gene × environment interaction in major depressive disorder. World J. Clin. Cases 9, 9368–9375. 10.12998/wjcc.v9.i31.9368.

110. Wilke, S.A., Johnson, C.L., Corlier, J., Marder, K.G., Wilson, A.C., Pleman, C.M., and Leuchter, A.F. (2022). Psychostimulant use and clinical outcome of repetitive transcranial magnetic stimulation treatment of major depressive disorder. Depress. Anxiety 39, 397–406. 10.1002/da.23255.

111. Hunter, A.M., Minzenberg, M.J., Cook, I.A., Krantz, D.E., Levitt, J.G., Rotstein, N.M., Chawla, S.A., and Leuchter, A.F. (2019). Concomitant medication use and clinical outcome of repetitive Transcranial Magnetic Stimulation (rTMS) treatment of Major Depressive Disorder. Brain Behav. 9, e01275. 10.1002/brb3.1275.

112. Cole, E., O’Sullivan, S.J., Tik, M., and Williams, N.R. (2024). Accelerated Theta Burst Stimulation: Safety, Efficacy, and Future Advancements. Biol. Psychiatry 95, 523–535. 10.1016/j.biopsych.2023.12.004.

113. Nandi, A., Virmani, G., Barve, A., and Marathe, S. (2021). DBscorer: An Open-Source Software for Automated Accurate Analysis of Rodent Behavior in Forced Swim Test and Tail Suspension Test. eNeuro 8, ENEURO.0305-21.2021. 10.1523/ENEURO.0305-21.2021.

114. Mathis, A., Mamidanna, P., Cury, K.M., Abe, T., Murthy, V.N., Mathis, M.W., and Bethge, M. (2018). DeepLabCut: markerless pose estimation of user-defined body parts with deep learning. Nat. Neurosci. 21, 1281–1289. 10.1038/s41593-018-0209-y.

115. Gabriel, C.J., Zeidler, Z., Jin, B., Guo, C., Goodpaster, C.M., Kashay, A.Q., Wu, A., Delaney, M., Cheung, J., DiFazio, L.E., et al. (2022). BehaviorDEPOT is a simple, flexible tool for automated behavioral detection based on markerless pose tracking. eLife 11, e74314. 10.7554/eLife.74314.

116. Schneider, C.A., Rasband, W.S., and Eliceiri, K.W. (2012). NIH Image to ImageJ: 25 years of image analysis. Nat. Methods 9, 671–675. 10.1038/nmeth.2089.

117. Arshadi, C., Günther, U., Eddison, M., Harrington, K.I.S., and Ferreira, T.A. (2021). SNT: a unifying toolbox for quantification of neuronal anatomy. Nat. Methods 18, 374–377. 10.1038/s41592-021-01105-7.

