## Supplemental Figures for "A cell type-specific mechanism driving the rapid antidepressant effects of transcranial magnetic stimulation"

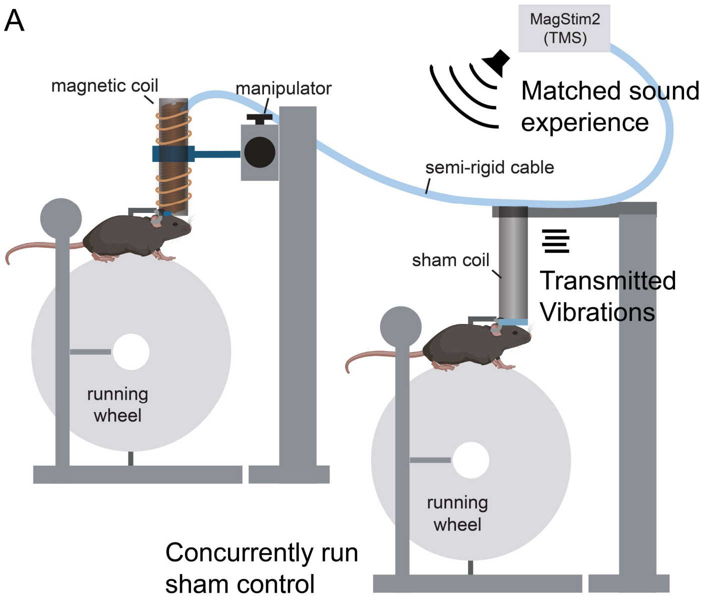


**Fig. S1.** Setup for concurrent aiTBS and sham treatment. (**A**) Sham-treated mice were run alongside experimental mice on an identical head-fixed setup with a sham coil pressed against their head. Sham mice experienced identical sounds from the stimulator. The semi-rigid cable connecting the TMS coil to the stimulator was draped over the sham coil to drive identically timed vibrations on the heads of the sham mice. Mouse diagrams were generated using Biorender.com.

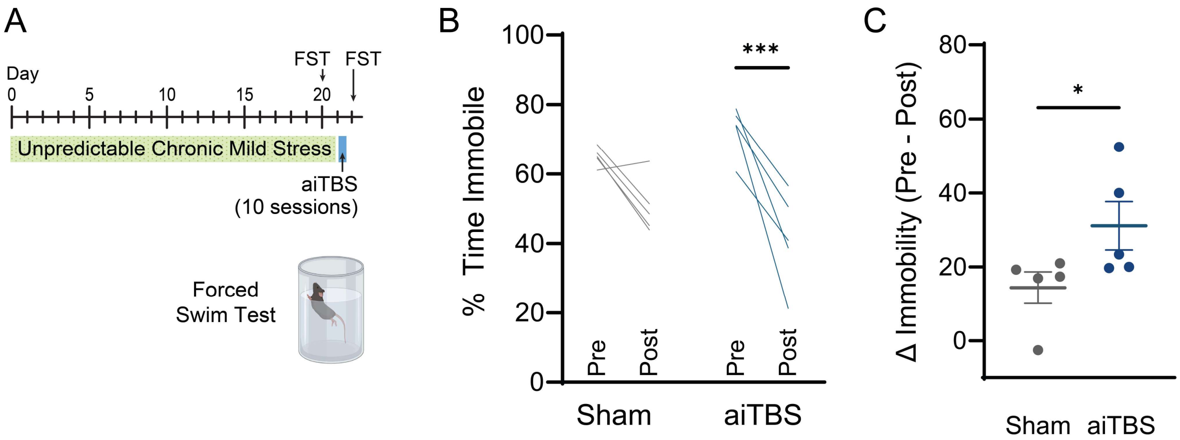


**Fig. S2. aiTBS reverses effects of unpredictable chronic mild stress on depression-related behavior.** (**A**) Experimental timeline. Mice underwent UCMS for 21 days, followed by one day of aiTBS treatment. FST was performed before and after aiTBS treatment. (**B**) Effects of aiTBS treatment on immobility in FST (Sham n=5, aiTBS n=5 mice; two-way RM ANOVA with Sidak's multiple comparisons test). (**C**) Change in immobility from pre- to post-treatment in aiTBS-treated or sham-treated mice (Sham n=5, aiTBS n=5 mice; unpaired t-test). For detailed statistical analysis see Table S1. All error bars reflect mean ± SEM. *$p<0.05$, ***$p<0.001$. Mouse diagrams were generated using Biorender.com.

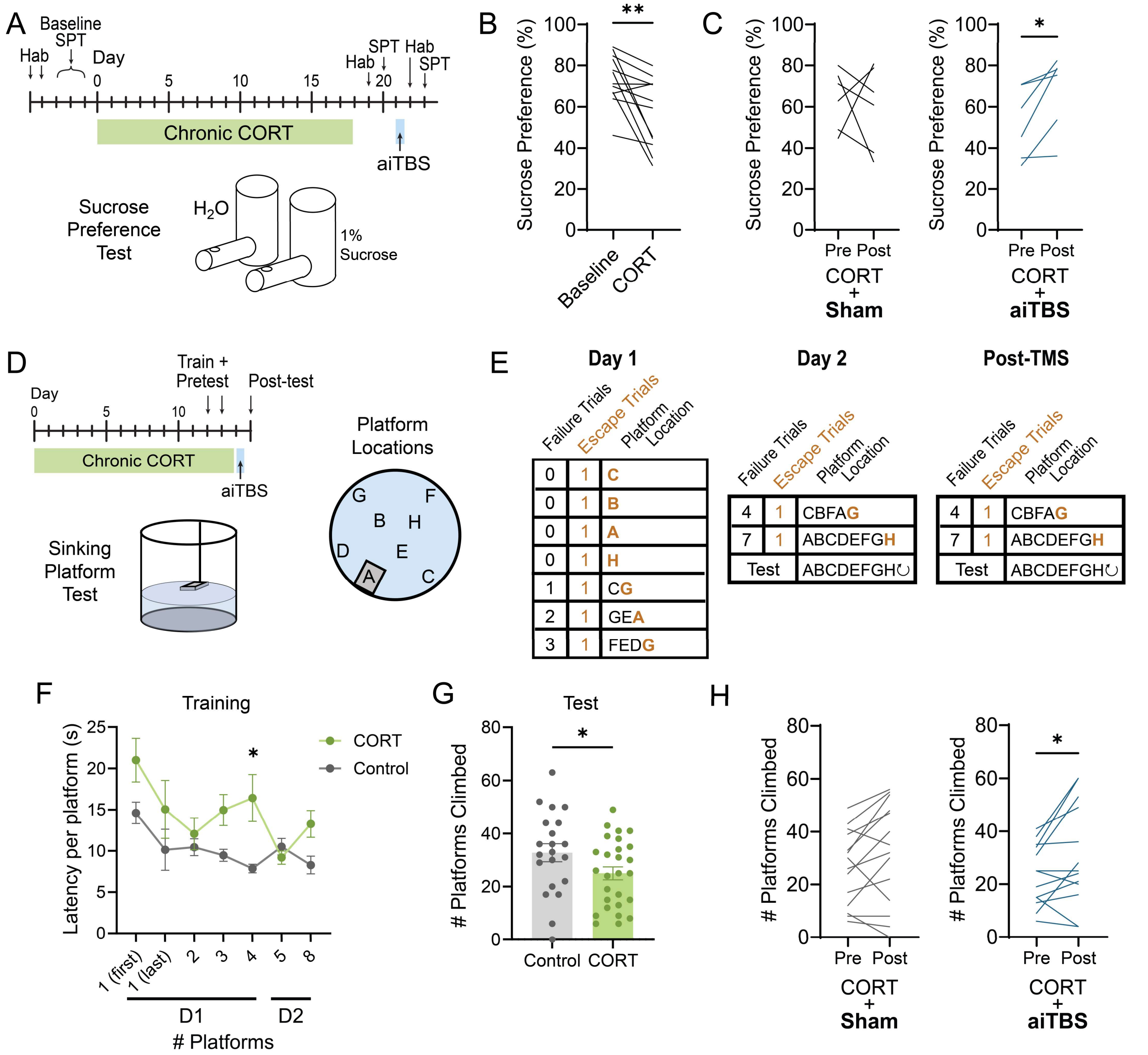


| Failure Trials | Escape Trials | Platform Location |
|---|---|---|
| 0 | 1 | C |
| 0 | 1 | B |
| 0 | 1 | A |
| 0 | 1 | H |
| 1 | 1 | CG |
| 2 | 1 | GEA |
| 3 | 1 | FEDG |

| Failure Trials | Escape Trials | Platform Location |
|---|---|---|
| 4 | 1 | CBFAG |
| 7 | 1 | ABCDEFGH |
| Test | | ABCDEFGH↻ |

| Failure Trials | Escape Trials | Platform Location |
|---|---|---|
| 4 | 1 | CBFAG |
| 7 | 1 | ABCDEFGH |
| Test | | ABCDEFGH↻ |

**Fig. S3. Effects of aiTBS on anhedonia and behavioral persistence.** (**A**) Experimental timeline for the sucrose preference test. Hab, habituation; SPT, sucrose preference test. (**B**) Effects of chronic CORT administration on sucrose preference (n=12 mice; paired student's t-test). (**C**) Effects of sham (left) or aiTBS treatment (right) on sucrose preference in mice having undergone chronic CORT (sham n=6, aiTBS n=6 mice; paired student's t-test). (**D**) Experimental timeline for the sinking platform test. In this assay, mice learn to climb a small platform to escape from a tank of water ("escape trial"). Then, "failure trials" are introduced, in which the experimenter sinks the platform after the mouse climbs it, requiring the mice to keep swimming. Mice must complete a progressively increasing number of failure trials before they are allowed an escape trial. Finally, during a test session with only failure trials, levels of persistence are quantified based on the total number of platforms climbed in a five-minute period. Letters indicate designated platform locations. (**E**) Training protocol. Orange letters indicate location for escape trials after the designated number of failure trials. (**F**) Average latency to climb the platform during training in non-stressed and CORT-treated mice. CORT-treated mice and non-stressed controls decreased their latency to climb platforms over time, but at the end of the first training day CORT-treated mice took longer to climb each platform (control n=21, CORT n=27 mice; two-way ANOVA with Sidak's multiple comparisons test). (**G**) Number of platforms climbed during the test session in non-stressed and CORT-treated mice. CORT-treated mice climbed fewer platforms than non-stressed controls (control n=21, CORT n=27 mice; unpaired t-test). (**H**) Effects of sham (left) or aiTBS treatment (right) on behavior in the test sessions before and after TMS. aiTBS- but not sham-treated mice climbed significantly more platforms compared to the pre-treatment baseline (Sham n=14, aiTBS n=13 mice; paired student's t-test). For detailed statistical analysis see Table S1. All error bars reflect mean ± SEM. *p<0.05, **p<0.01.

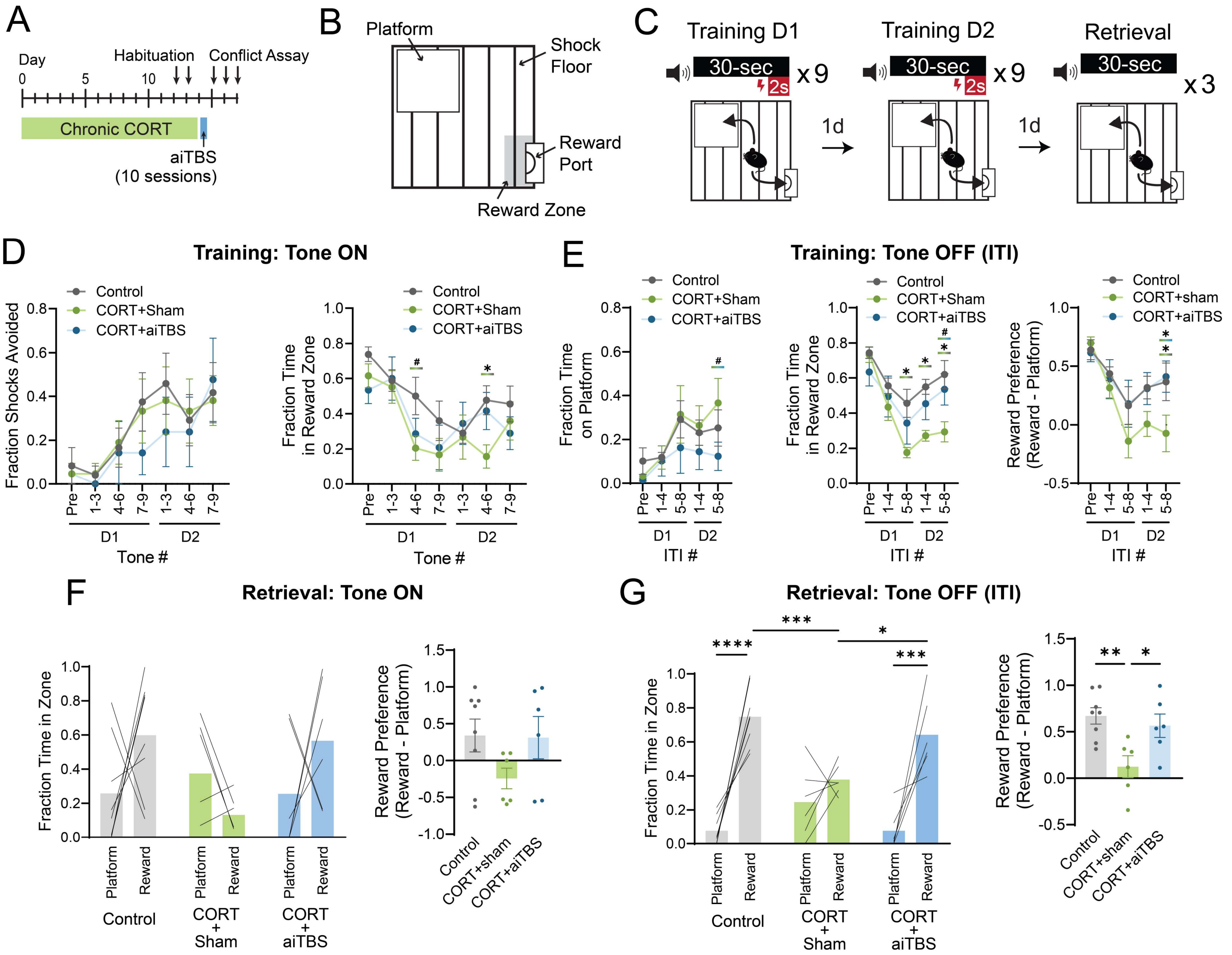


**Fig. S4. Effects of aiTBS on behavior in an approach-avoidance conflict assay.** (**A**,**B**,**C**) Timeline, chamber design and protocol for the conflict assay. In this assay, mice can access a reward port that delivers sweetened condensed milk, but at random times, they are presented with a tone that co-terminates with a mild foot shock. Mice learn they can avoid the shock by navigating to a safety platform in the opposite corner, where rewards are inaccessible. Mice were exposed to nine tone-shock pairings on days 1 and 2. On day 2, they were exposed to three tones during a tone-only retrieval session to examine memory-guided behavior. (**D**) Summaries of fraction of shocks avoided (left) and fraction time spent in reward zone (right) during the tone ON period on training days (D) 1 and 2 (non-stressed control n=8, CORT+Sham n=7, CORT+aiTBS n=7 mice; repeated measures two-way ANOVA with Tukey's multiple comparisons test). (**E**) Summaries of fraction of shocks avoided (left), fraction time spent in reward zone (center), and reward zone preference (time in reward zone-time in platform zone) during Tone OFF periods (inter-tone interval, ITI) (non-stressed control n=8, CORT+Sham n=7, CORT+aiTBS n=7 mice; repeated measures two-way ANOVA with Tukey's multiple comparisons test). (**F**) Summaries of fraction time in platform and reward zones (left) and reward preferences (right) during Retrieval Tone ON periods (non-stressed control n=8, CORT+Sham n=6, CORT+aiTBS n=6 mice; Left, two-way RM ANOVA with Tukey's multiple comparisons test; Right, one-way ANOVA with Tukey's multiple comparisons test). (**G**) Summaries of fraction time in platform and reward zones (left) and reward preferences (right) during Retrieval Tone OFF (ITI) periods (non-stressed control n=8, CORT+Sham n=6, CORT+aiTBS n=6 mice; Left, two-way RM ANOVA with Sidak's multiple comparisons test; Right, one-way ANOVA with Tukey's multiple comparisons test). For detailed statistical analysis see Table S1. All error bars reflect mean ± SEM. #p<0.1, *p<0.05, **p<0.01, ***p<0.001, ****p<0.0001.

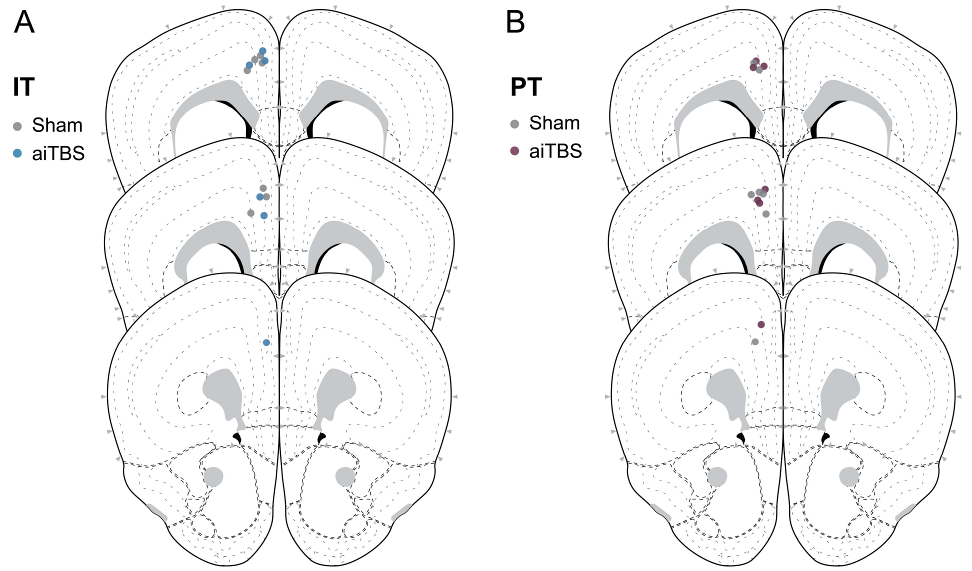


**Fig. S5. Fiber targeting for fiber photometry experiments.** (**A**) Locations of fiber tips for recordings of IT neurons. Sham animals are shown in gray and aiTBS animals are shown in blue. (**B**) Locations of fiber tips for recordings of PT neurons. Sham animals are shown in gray and aiTBS animals are shown in purple.

---

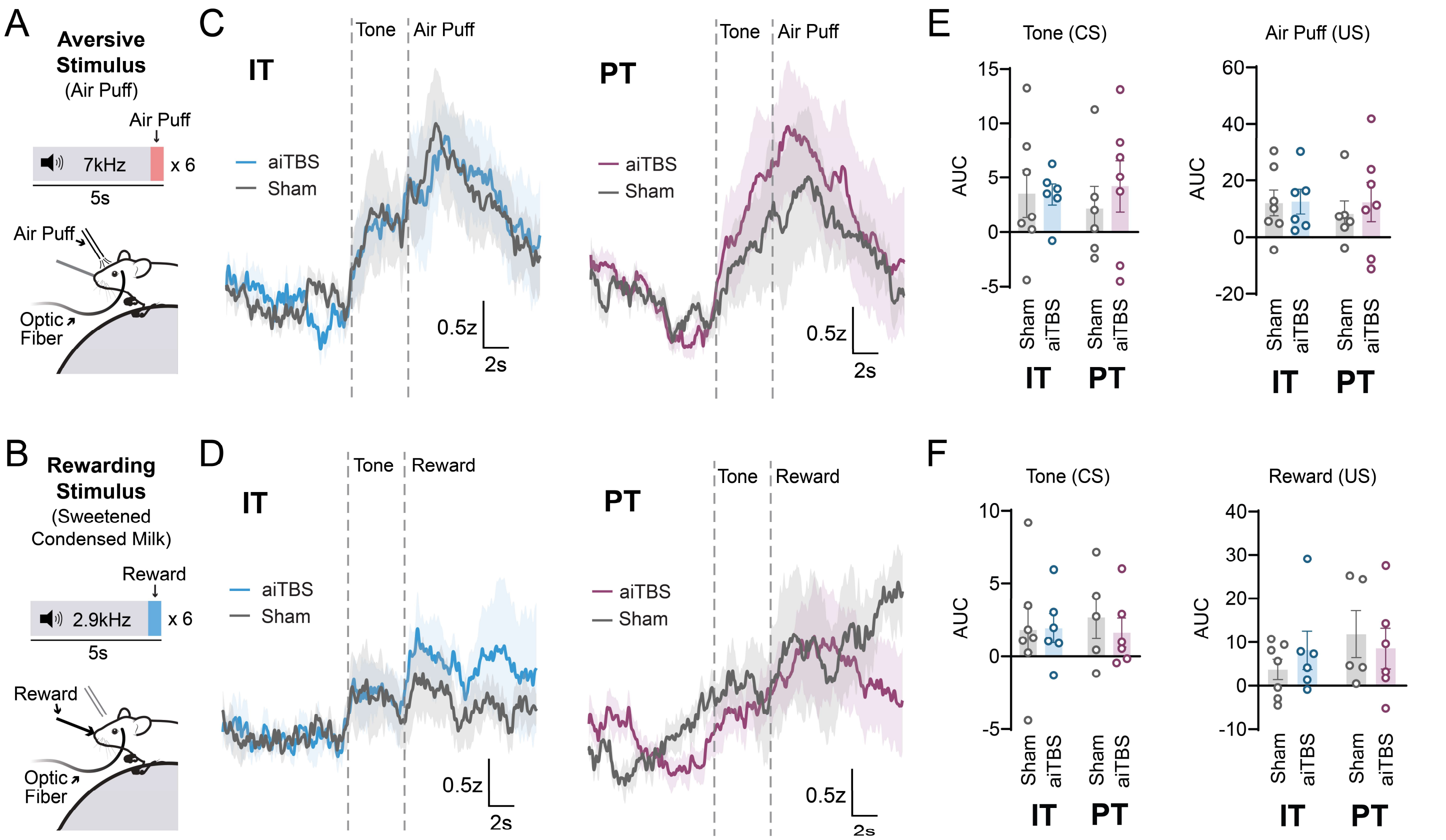


**Fig. S6. Fiber photometry recording from aversive and rewarding Pavlovian stimuli following aiTBS treatment.** Experimental setup for recording neural responses to aversive and rewarding stimuli. Mice were presented with randomly interspersed 5s tones at 7kHz and 2.9kHz co-terminating with air puff (**A**) or reward delivery (**B**), respectively. (**C,D**) Fiber photometry recordings from IT and PT neurons in this assay following aiTBS or sham treatment. (**E,F**) AUC from conditioned and unconditioned aversive and rewarding stimuli (IT sham n=7, IT aiTBS n=6, PT sham n=6, PT aiTBS n=7 mice; two-way ANOVA with Sidak's multiple comparisons test). For detailed statistical analysis see Table S1. Lines and shading reflect mean ± SEM of average traces across mice. All error bars reflect mean ± SEM.

---

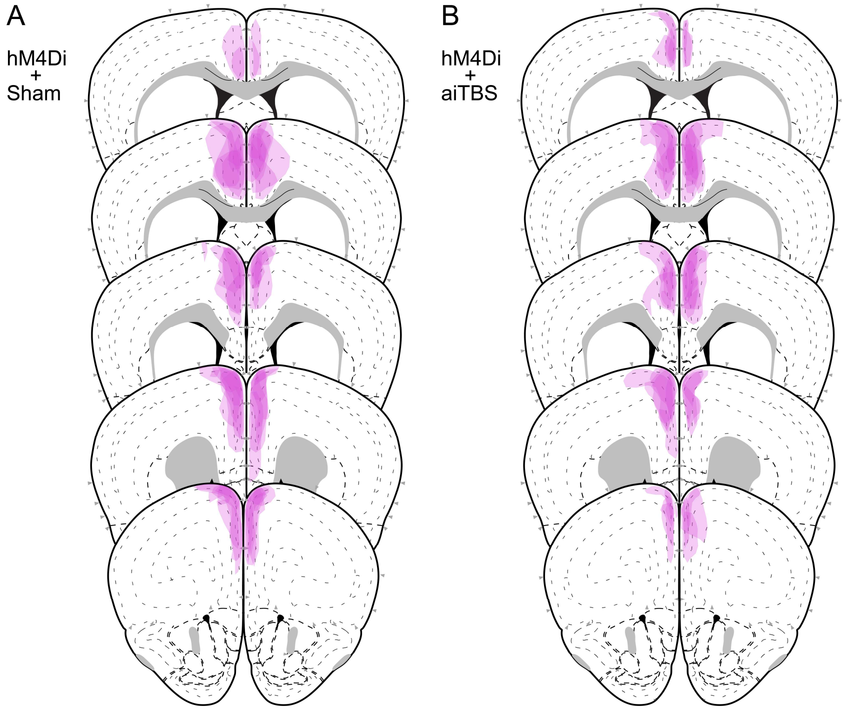


**Fig. S7. Viral targeting and hM4Di expression in dmPFC IT neurons** in sham-treated (**A**) and aiTBS-treated animals (**B**). Expression for each animal is demonstrated by a translucent outline.

**Table S1.** Detailed statistical analyses.

| Figure | Sample Size | Test | Result | |
|---|---|---|---|---|
| Figure 1E | n=12 Control<br>n=24 UCMS<br>n=15 CORT | One-way ANOVA with Tukey's Multiple Comparisons | F (2, 48) = 14.01 | P<0.0001 |
| Figure 1F | n=8 Control<br>n=8 CORT | Unpaired t-test (one-tailed) | t=2.703, df=14 | P=0.0086 |
| Figure 1G | n=8 Control<br>n=8 CORT | Unpaired t-test (one-tailed) | t=0.5199, df=14 | P=0.6944 |
| Figure 1I | n=11 CORT + Sham<br>n=10 CORT + aiTBS | Welch's t-test (one-tailed) | t=2.177, df=12.38 | P=0.0247 |
| Figure 1J | n=11 CORT + Sham<br>n=10 CORT + aiTBS | Unpaired t-test (one-tailed) | t=1.591, df=19 | P=0.0641 |
| Figure 1K | n=11 CORT + Sham<br>n=10 CORT + aiTBS | Welch's t-test (one-tailed) | t=1.765, df=10.88 | P=0.0528 |
| Figure 1L | n=11 CORT + Sham<br>n=10 CORT + aiTBS | Welch's t-test (one-tailed) | t=3.381, df=10.52 | P=0.0033 |
| Figure 2F, left | n=6 IT sham<br>n=5 IT aiTBS<br>n = 7 PT sham<br>n = 8 PT aiTBS | Two-Way ANOVA with Sidak's Multiple Comparisons Test | Cell Type: F (1, 22) = 0.2240<br>Treatment: F (1, 22) = 11.44<br>Interaction: F (1, 22) = 0.2240 | P=0.6407<br>P=0.0027<br>P=0.6407 |
| Figure 2F, right | n=6 IT sham<br>n=5 IT aiTBS<br>n = 7 PT sham<br>n = 8 PT aiTBS | Two-Way ANOVA with Sidak's Multiple Comparisons Test | Cell Type: F (1, 22) = 3.498<br>Treatment: F (1, 22) = 6.935<br>Interaction: F (1, 22) = 3.498 | P=0.0748<br>P=0.0152<br>P=0.0748 |
| Figure 2H | n=6 IT sham<br>n=5 IT aiTBS<br>n = 7 PT sham<br>n = 8 PT aiTBS | Two-Way RM ANOVA with Sidak's Multiple Comparisons Test | Cell Type: F (1, 11) = 12.48<br>Time: F (5, 55) = 2.204<br>Interaction: F (5, 55) = 6.083 | P=0.0047<br>P=0.0669<br>P=0.0001 |
| Figure 3A | n=13 Sham<br>n=13 aiTBS | Unpaired t-test (one-tailed) | t=1.336, df=24 | P=0.0971 |

| | | | | |
|---|---|---|---|---|
| Figure 3D | n=7 IT sham<br>n=6 IT aiTBS<br>n = 5 PT sham<br>n = 7 PT aiTBS | Two-Way ANOVA with Sidak's Multiple Comparisons Test | Cell Type: F (1, 21) = 2.985<br>Treatment: F (1, 21) = 1.318<br>Interaction: F (1, 21) = 5.686 | P=0.0987<br>P=0.2639<br>P=0.0266 |
| Figure 3E | n=7 IT sham<br>n=6 IT aiTBS<br>n = 5 PT sham<br>n = 7 PT aiTBS | Two-Way ANOVA with Sidak's Multiple Comparisons Test | Cell Type: F (1, 21) = 0.03385<br>Treatment: F (1, 21) = 3.419<br>Interaction: F (1, 21) = 3.032 | P=0.8558<br>P=0.0786<br>P=0.0963 |
| Figure 3F | n=7 IT sham<br>n=6 IT aiTBS<br>n = 5 PT sham<br>n = 7 PT aiTBS | Two-Way ANOVA with Sidak's Multiple Comparisons Test | Cell Type: F (1, 21) = 7.967<br>Treatment: F (1, 21) = 0.9558<br>Interaction: F (1, 21) = 5.929 | P=0.0102<br>P=0.3394<br>P=0.0239 |
| Figure 3H | n=7 IT sham<br>n=6 IT aiTBS<br>n = 6 PT sham<br>n = 6 PT aiTBS | Two-Way ANOVA with Sidak's Multiple Comparisons Test | Cell Type: F (1, 21) = 1.363<br>Treatment: F (1, 21) = 2.516<br>Interaction: F (1, 21) = 4.173 | P=0.2561<br>P=0.1276<br>P=0.0538 |
| Figure 3J | n=7 IT sham<br>n=6 IT aiTBS<br>n = 6 PT sham<br>n = 6 PT aiTBS | Two-Way ANOVA with Sidak's Multiple Comparisons Test | Cell Type: F (1, 21) = 0.8493<br>Treatment: F (1, 21) = 2.339<br>Interaction: F (1, 21) = 4.894 | P=0.3672<br>P=0.1411<br>P=0.0382 |
| Figure 4E | n=3 naïve<br>n=3 CORT + sham<br>n=3 CORT + aiTBS | Two-Way RM ANOVA with Sidak's Multiple Comparisons Test | Dendritic Segment: F (1, 6) = 3.475<br>Treatment: F (2, 6) = 26.74<br>Interaction: F (2, 6) = 1.041<br>Subject: F (6, 6) = 2.218 | P=0.1116<br>P=0.0010<br>P=0.4090<br>P=0.1776 |
| Figure 4G | n=3 naïve<br>n=4 CORT + sham<br>n=4 CORT + aiTBS | Two-Way RM ANOVA with Sidak's Multiple Comparisons Test | Dendritic Segment: F (1, 8) = 23.67<br>Treatment: F (2, 8) = 8.851<br>Interaction: F (2, 8) = 5.055<br>Subject: F (8, 8) = 0.9515 | P=0.0012<br>P=0.0094<br>P=0.0381<br>P=0.5272 |
| Figure 5B | n=3 mCherry + aiTBS<br>n=4 hM4Di + aiTBS | Unpaired t-test (one-tailed) | t=15.29, df=5 | P<0.0001 |

| | | | | |
|---|---|---|---|---|
| Figure 5D | n=7 mCherry + sham<br>n=6 mCherry + aiTBS<br>n=5 hM4Di + sham<br>n=6 hM4Di + aiTBS | Three-Way RM ANOVA with Sidak's Multiple Comparisons Test | Pre/Post: F (1, 20) = 23.43<br>Virus: F (1, 20) = 1.444<br>Treatment: F (1, 20) = 0.5375<br>Pre/Post x Virus: F (1, 20) = 6.621<br>Pre/Post x Treatment: F (1, 20) = 1.527<br>Virus x Treatment: F (1, 20) = 0.9805<br>Pre/Post x Virus x Treatment: F (1, 20) = 2.472 | P<0.0001<br>P=0.2435<br>P=0.4720<br>P=0.0182<br>P=0.2309<br>P=0.3339<br>P=0.1316 |
| Figure 5E | n=6 mCherry + aiTBS<br>n=6 hM4Di + aiTBS | Unpaired t-test (two-tailed) | t=2.831, df=10 | P=0.0178 |
| Figure S2B | n=5 sham<br>n=5 aiTBS | Two-Way RM ANOVA with Sidak's Multiple Comparisons Test | Condition: F (1, 8) = 0.02118<br>Pre/Post: F (1, 8) = 34.10<br>Interaction: F (1, 8) = 4.584<br>Subject: F (8, 8) = 0.9819 | P=0.8879<br>P=0.0004<br>P=0.0647<br>P=0.5100 |
| Figure S2C | n=5 sham<br>n=5 aiTBS | Unpaired t-test (one-tailed) | t=2.141, df=8 | P=0.0323 |
| Figure S3B | n=12 mice | Paired t-test (two-tailed) | t=3.199, df=11 | P=0.0085 |
| Figure S3C | n=6 sham<br>n=6 aiTBS | Paired t-test (two-tailed) | sham: t=0.3644, df=5<br>aiTBS: t=2.917, df=5 | P=0.7305<br>P=0.0331 |
| Figure S3F | n=21 Control<br>n=27 CORT | Two-Way RM ANOVA with Sidak's Multiple Comparisons Test | Stress: F (1, 46) = 4.832<br>Trial: F (6, 276) = 5.258<br>Interaction: F (6, 276) = 2.099<br>Subject: F (46, 276) = 5.508 | P=0.0330<br>P<0.0001<br>P=0.0536<br>P<0.0001 |
| Figure S3G | n=21 Control<br>n=27 CORT | Unpaired t-test (one-tailed) | t=1.899, df=46 | P=0.0319 |
| Figure S3H | n=14 sham<br>n=13 aiTBS | Paired t-test (two-tailed) | sham: t=1.847, df=13<br>aiTBS: t=2.251, df=12 | P=0.0876<br>P=0.0440 |
| Figure S4D, left | n=8 Control<br>n=7 CORT+Sham<br>n=7 CORT+aiTBS | Two-Way RM ANOVA with Tukey's Multiple Comparisons Test | Condition: F (2, 19) = 0.2462<br>Tone bin: F (6, 114) = 7.898<br>Interaction: F (12, 114) = 0.4337<br>Subject: F (19, 114) = 5.662 | P=0.7842<br>P<0.0001<br>P=0.9468<br>P<0.0001 |

| | | | | |
|---|---|---|---|---|
| Figure S4D, right | n=8 Control<br>n=7 CORT+Sham<br>n=7 CORT+aiTBS | Two-Way RM ANOVA with Tukey's Multiple Comparisons Test | Condition: F (2, 19) = 1.793<br>Tone bin: F (6, 114) = 10.51<br>Interaction: F (12, 114) = 1.060<br>Subject: F (19, 114) = 4.190 | P=0.0935<br>P<0.0001<br>P=0.4002<br>P<0.0001 |
| Figure S4E, left | n=8 Control<br>n=7 CORT+Sham<br>n=7 CORT+aiTBS | Two-Way RM ANOVA with Tukey's Multiple Comparisons Test | Condition: F (2, 19) = 0.8987<br>ITI bin: F (4, 76) = 8.649<br>Interaction: F (8, 76) = 0.8933<br>Subject: F (19, 76) = 6.398 | P=0.5264<br>P<0.0001<br>P=0.4237<br>P<0.0001 |
| Figure S4E, center | n=8 Control<br>n=7 CORT+Sham<br>n=7 CORT+aiTBS | Two-Way RM ANOVA with Tukey's Multiple Comparisons Test | Condition: F (2, 19) = 3.706<br>ITI bin: F F (4, 76) = 19.54<br>Interaction: F (8, 76) = 1.982<br>Subject: F F (19, 76) = 4.800 | P=0.0438<br>P<0.0001<br>P=0.0600<br>P<0.0001 |
| Figure S4E, right | n=8 Control<br>n=7 CORT+Sham<br>n=7 CORT+aiTBS | Two-Way RM ANOVA with Tukey's Multiple Comparisons Test | Condition: F (2, 19) = 1.617<br>ITI bin: F (4, 76) = 17.87<br>Interaction: F (8, 76) = 1.734<br>Subject: F (19, 76) = 5.975 | P=0.1041<br>P<0.0001<br>P=0.2246<br>P<0.0001 |
| Figure S4F, left | n=8 Control<br>n=6 CORT+Sham<br>n=6 CORT+aiTBS | Two-Way RM ANOVA with Tukey's Multiple Comparisons Test | Condition: F (2, 17) = 5.264<br>Location: F (1, 17) = 1.061<br>Interaction: F (2, 17) = 1.997<br>Subject: F (17, 17) = 0.1311 | P=0.0166<br>P=0.3175<br>P=0.1663<br>P>0.9999 |
| Figure S4F, right | n=8 Control<br>n=6 CORT+Sham<br>n=6 CORT+aiTBS | One-way ANOVA with Tukey's Multiple Comparisons | F (2,17) = 1.997 | P=0.1663 |
| Figure S4G, left | n=8 Control<br>n=6 CORT+Sham<br>n=6 CORT+aiTBS | Two-Way RM ANOVA with Tukey's Multiple Comparisons Test | Condition: F (2, 17) = 2.737<br>Location: F (1, 17) = 51.39<br>Interaction:F (2, 17) = 6.960<br>Subject:F (17, 17) = 0.3021 | P=0.0932<br>P<0.0001<br>P=0.0062<br>P=0.9910 |
| Figure S4G, right | n=8 Control<br>n=6 CORT+Sham<br>n=6 CORT+aiTBS | One-way ANOVA with Tukey's Multiple Comparisons | F (2,17) = 6.960 | P=0.0062 |
| Figure S6C, left | n=7 IT sham<br>n=6 IT aiTBS<br>n = 6 PT sham<br>n = 7 PT aiTBS | Two-Way ANOVA with Sidak's Multiple Comparisons Test | Cell Type: F (1, 22) = 0.02126<br>Treatment: F (1, 22) = 0.2313<br>Interaction: F (1, 22) = 0.2747 | P=0.8854<br>P=0.6353<br>P=0.6054 |
| Figure S6C, right | n=7 IT sham<br>n=6 IT aiTBS<br>n = 6 PT sham<br>n = 7 PT aiTBS | Two-Way ANOVA with Sidak's Multiple Comparisons Test | Cell Type: F (1, 22) = 0.1367<br>Treatment: F (1, 22) = 0.1850<br>Interaction: F (1, 22) = 0.1143 | P=0.7151<br>P=0.6713<br>P=0.7385 |

| | | | | |
|---|---|---|---|---|
| Figure S6F, left | n=7 IT sham<br>n=6 IT aiTBS<br>n = 5 PT sham<br>n = 6 PT aiTBS | Two-Way ANOVA with Sidak's Multiple Comparisons Test | Cell Type: $F (1, 20) = 0.1233$<br>Treatment: $F (1, 20) = 0.1921$<br>Interaction: $F (1, 20) = 0.04358$ | $P=0.8367$<br>$P=0.7291$<br>$P=0.6659$ |
| Figure S6F, right | n=7 IT sham<br>n=6 IT aiTBS<br>n = 5 PT sham<br>n = 6 PT aiTBS | Two-Way ANOVA with Sidak's Multiple Comparisons Test | Cell Type: $F (1, 20) = 1.041$<br>Treatment: $F (1, 20) = 0.01959$<br>Interaction: $F (1, 20) = 0.8549$ | $P=0.3197$<br>$P=0.8901$<br>$P=0.3662$ |
